# Identification of a forkhead box protein transcriptional network induced in human neutrophils in response to inflammatory stimuli

**DOI:** 10.1101/2022.12.02.518811

**Authors:** Aiten Ismailova, Reyhaneh Salehi-Tabar, Vassil Dimitrov, Babak Memari, Camille Barbier, John H. White

## Abstract

Neutrophils represent the largest proportion of circulating leukocytes and, in response to inflammatory stimuli, are rapidly recruited to sites of infection where they neutralize pathogens. We have identified a novel neutrophil transcription network induced in response to inflammatory stimuli. We performed the first RNAseq analysis of human neutrophils exposed to lipopolysaccharide (LPS), followed by a meta-analysis of our dataset and previously published studies of LPS-challenged neutrophils. This revealed a robustly enhanced transcriptional network driven by forkhead box (FOX) transcription factors. The network is enriched in genes encoding proinflammatory cytokines and transcription factors, including *MAFF* and *ATF3*, which are implicated in responses to stress, survival and inflammation. Expression of transcription factors FOXP1 and FOXP4 is induced in neutrophils exposed to inflammatory stimuli, and potential FOXP1/FOXP4 binding sites were identified in several genes in the network, all located in chromatin regions consistent with neutrophil enhancer function. Chromatin immunoprecipitation (ChIP) assays in neutrophils confirmed enhanced binding of FOXP4, but not FOXP1, to multiple sites in response to LPS. Binding to numerous motifs and transactivation of network genes were also observed when FOXP proteins were transiently expressed in HEK293 cells. In addition to LPS, the transcriptional network is induced by other inflammatory stimuli, indicating it represents a general neutrophil response to inflammation. Collectively, these findings reveal a role for the FOXP4 transcription network as a regulator of responses to inflammatory stimuli in neutrophils.

**Author Summary:** In response to pathogens, neutrophils, the most abundant white blood cells in the body, are the first to be recruited to sites of infection. However, defects in neutrophil responses lead to common chronic inflammatory conditions such as atherosclerosis, chronic obstructive lung disease and autoimmune disorders. As such, it is critical to uncover the molecular players implicated in neutrophil responses to signals that induce inflammation. Here we profile how bacterial lipopolysaccharide (LPS), which is derived from the cell walls of bacteria and is a commonly used agent to mimic inflammation, alters gene transcription in isolated human neutrophils. We also combined our data with those of other published studies to identify conserved molecular pathways stimulated in LPS-exposed neutrophils. This analysis revealed a network of genes whose transcription is regulated by members of the so-called forkhead box (FOX) transcription factors. We provide evidence that FOXP4 regulates transcription of genes within the network in neutrophils. We also find that the same network of genes is induced by other inflammatory stimuli, suggesting it plays a role in neutrophil responses to inflammation.

## Introduction

Neutrophils represent the largest proportion of circulating leukocytes in the blood and are essential to the innate immune response against invading pathogens (1). In response to inflammatory stimuli, neutrophils are rapidly recruited to sites of infection and inflammation where they efficiently bind, engulf and inactivate pathogens. Moreover, these cells secrete chemokines and pro-inflammatory cytokines, which facilitates recruitment and activation of additional neutrophils as well as other immune cells to inflamed tissues (1). Defects in neutrophil antimicrobial processes or a decrease in neutrophil abundance often lead to increased risk of infection or chronic inflammatory or autoimmune conditions such as systemic lupus erythematosus and rheumatoid arthritis (2–4). They are also implicated in the pathogenesis of viral infections; for example, a growing body of evidence supports a link between infiltrating activated neutrophils and increased COVID-19 disease severity (5–7). The killing of invading microorganisms by neutrophils thus necessitates careful control of neutrophil function, as neutropenia renders the body vulnerable to infection, whereas overactivity is associated with inflammatory diseases (1). Moreover, neutrophils can influence cancer progression; they are abundant in tumours and promote tumour development by secreting cytokines and matrix-degrading proteases (2). Consequently, a comprehensive understanding of (dys)regulated molecular signal pathways and transcriptional networks in activated neutrophils is critical for understanding responses to infectious or inflammatory signals.

Bacterial lipopolysaccharide (LPS) is often used as an inflammatory signal, and is capable of inducing several functional responses in neutrophils that contribute to innate immunity, such as altering neutrophil adhesion, respiratory burst, degranulation and motility (8). It follows that transcriptional responses play a key role in the stimulation of cytokine and chemokine production in neutrophils. LPS signaling activates transcription factors, such as nuclear factor-kappa B (NF-κB) and STAT3 (9, 10). Several microarray studies of varying sizes have screened human neutrophil LPS-mediated signal transduction pathways and transcriptional networks (11–19). Despite discrepancies in microarray platforms and doses of LPS treatment, several studies have shown that transcripts encoding the pro-inflammatory cytokines and chemokines, IL-1B, IL-6, CCL2, and CCL4 are induced in response to the inflammatory agent.

Here, we performed the first RNA sequencing (RNAseq) analysis of primary human neutrophils to probe the transcriptional responses to LPS on a genome-wide scale, followed by a meta-analysis of LPS-induced neutrophil gene expression profiles. We found that the most robustly enhanced transcriptional network, which is driven by forkhead box transcription factors, consists of genes that encode pro-inflammatory cytokines and transcription factors that are associated with inflammation and stress, such as MAFF and ATF3. Forkhead box transcription factors are characterized by the presence of a highly conserved forkhead DNA-binding domain (20). They possess overlapping binding specificities and are expressed in a cell-specific manner (21). We find that, among forkhead transcription factors, expression of FOXP1 and FOXP4 is enhanced in primary human neutrophils in response to LPS. This leads to enhanced binding of FOXP4, but not FOXP1, to multiple forkhead box motifs in network genes in neutrophils. Moreover, the network is induced in neutrophils by other inflammatory signals, such as a PKA agonist, GM-CSF and IFN-γ, indicating that it is a component of neutrophil responses to inflammation. These studies reveal a novel transcription network induced in neutrophils in response to inflammatory stimuli.

## Results

### Gene expression profiling by RNAseq of primary human neutrophils stimulated with LPS

To determine the transcriptomic responses of neutrophils under inflammatory conditions, we performed gene expression profiling by RNAseq in triplicate isolates of primary human neutrophils treated with LPS or vehicle (Fig 1A, S1 File). Cells were stimulated with 1µg/ml LPS or treated with vehicle for 6 hours to gain insight into primary and secondary effects on gene expression. 1 µg/ml of LPS was employed in order to mimic systemic inflammation; this dose was shown to induce the greatest ERK phosphorylation in human neutrophils (28). ERK is a subfamily of the mitogen-activated protein kinase (MAPK), which, upon phosphorylation, is involved in signal transduction pathways of inflammation (29, 30) and neutrophil phagocytosis (31). Moreover, 1 µg/ml of LPS was found to be efficient at halting primary human neutrophil chemotaxis at infectious foci, enabling the cells to exert bactericidal functions at pathogenic sites (32). LPS treatment resulted in broad changes in mRNA profiles, in which 2565 and 2457 genes were up- and downregulated, respectively, at least 2-fold or greater compared with vehicle (p≤0.05). We then examined the cell signaling canonical pathways, diseases and biological functions as well as upstream transcriptional regulators by pathways analysis (Fig 1A). Up- and downregulated genes with a range of fold regulations were subsequently selected for RT/qPCR validation of the RNAseq analysis. Results from RT/qPCR gene expression analysis largely corroborate the expression data generated by RNAseq (Fig 1B). As we used a higher dose of LPS than other studies, we evaluated the effects of differing doses on neutrophil gene expression. Dose-response curves were generated by exposing neutrophils to increasing concentrations of LPS (0, 0.01, 0,1, 1 µg/ml) for 6 h and assessing the regulation of validated genes by RT/qPCR (Fig 1C). This revealed that the higher dose used in our study did not differ substantially from lower doses in its effects on gene repression (*CAMP, CXCR2*), but appears to have been less efficacious in activating gene expression (*IL36G, CSF3, CXCL8*). Importantly, however, these data suggest that it is unlikely that the higher dose used here led to gene regulatory events that would be absent in other studies that employed lower doses of LPS.

**Fig 1.**
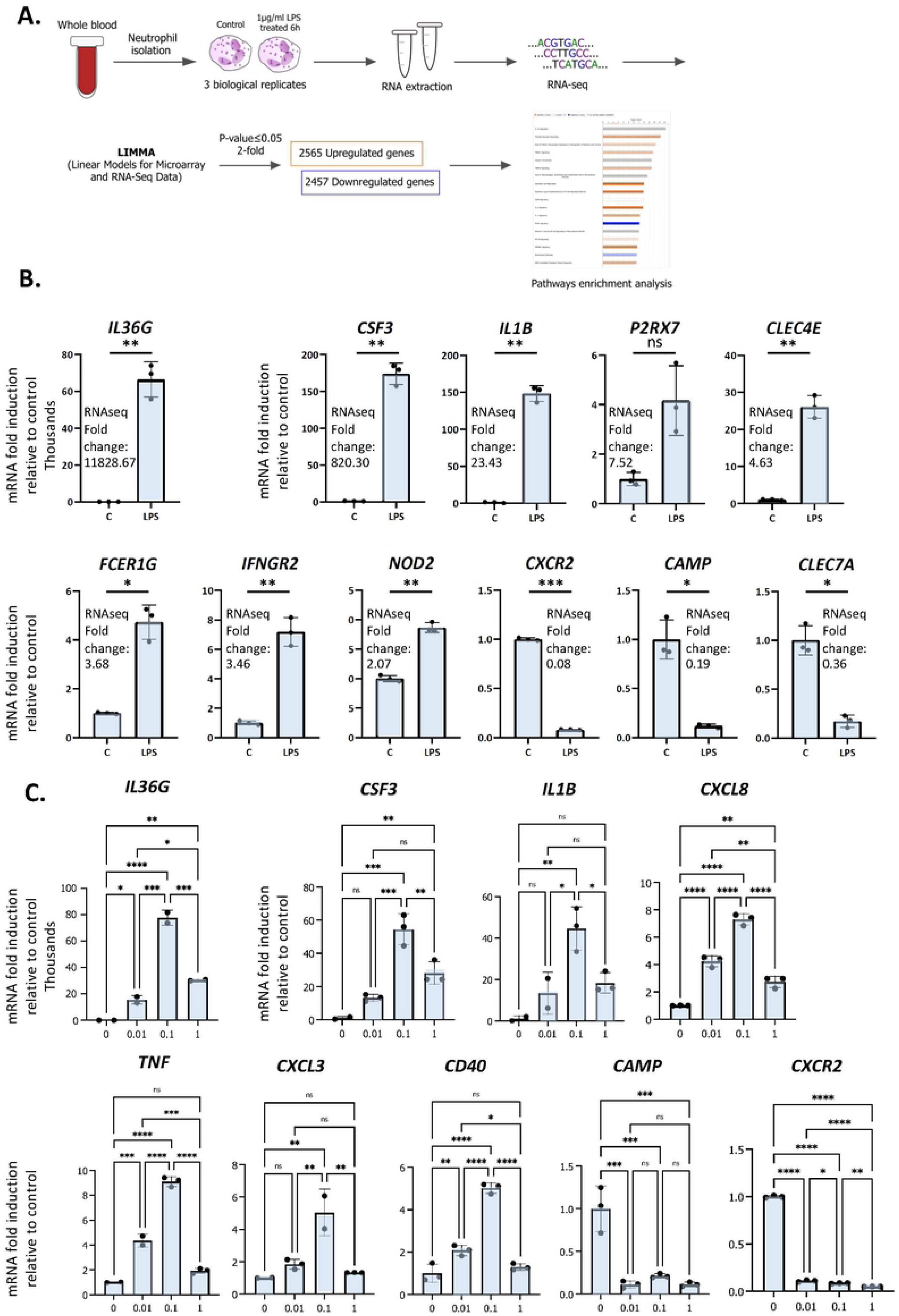
RNAseq analysis of primary human neutrophils. **A.** Schematic representation of workflow for RNAseq analysis of LPS-regulated gene expression in neutrophils. **B.** Validation of regulation by LPS of differentially expressed genes by RT/qPCR analysis. Data are representative of 2 or 3 biological replicates. *P ≤ 0.05, **P ≤ 0.01, ***P ≤ 0.001 and ns ≥0.05 as determined by paired Student’s t-test for technical replicates of one representative sample. **C.** Dose-response curves of LPS. Primary human neutrophils were treated for 6h with 0 to 1 µg of LPS per ml and RT/qPCR was subsequently performed. Graphics are mean ± SD from 3 technical replicates and data are representative of 2 or 3 biological replicates. *P ≤ 0.05, **P ≤ 0.01, ***P ≤ 0.001, ****P ≤ 0.0001 and ns ≥0.05 as determined by one-way ANOVAs followed by Tukey’s post hoc test for multiple comparisons.

### Meta-analysis of LPS-challenged neutrophils and identification of a forkhead box transcription factor network

To compare our data with those of other gene expression studies of LPS-stimulated human and mouse neutrophils, we conducted a meta-analysis of our RNAseq data and other available microarray and RNAseq expression profiles. Datasets from the Gene Expression Omnibus (GEO) public repository with available raw data were included, which produced 14 human microarray studies and 2 mouse RNAseq expression profiles (Table S1). Differential gene expression analysis was carried out using the same pipeline to ensure that results were comparable (see Methods); the Linear Models for Microarray Data (LIMMA) Bioconductor package was used, as it performs well in different settings for microarray and RNAseq experiments (33). Quality control (QC) of RNAseq and microarray datasets was determined, and Figs. S3 and S4 provide examples of QC criteria for human and mouse expression profiles (see also materials and methods for details). We included profiles generated with neutrophils stimulated *in vitro* with different concentrations of LPS over time periods ranging from 1 to 16h (see Table S1 for details). *In vivo* profiles were derived from patients administered an intravenous bolus injection of LPS for 1-6h (human endotoxemia model), followed by isolation of cells from serum, either as total neutrophil populations or as subpopulations sorted for CD16 and CD62L expression levels, markers of neutrophil maturity (14, 17, 18). Generally, the *in vivo* profiles gave rise to much smaller and more heterogeneous datasets. Expression patterns of the sorted subpopulations and total neutrophils were essentially completely different (Fig S2), with the exception of partial overlap between total and CD16^dim^/CD62L^bright^ cells, a subset that arises rapidly upon experimental human endotoxemia (18, 34).

Our study represents the first RNAseq transcriptional analysis of LPS-stimulated human neutrophils and produced the largest number of up- and downregulated genes. Venn diagrams depicting up- and downregulated genes at least 2-fold in our data and other human *in vitro* and *in vivo* expression profiles from LPS-stimulated neutrophils revealed substantial overlap with other *in vitro* datasets (Figs. 2A, S1). Our data showed more enrichment than others for diseases and functions, such as chemotaxis, cell binding, viral infection and apoptosis, largely because of the greater number of regulated genes identified (Fig 2B). Notably, the canonical pathways regulated as well as upstream regulators identified in our profile were most similar to those of the smaller study of Khaenam et al (19), who also treated neutrophils *in vitro* for 6h with LPS (Figs. 2C, 3A). There was substantially less overlap in our regulated gene sets with those identified in *in vivo* studies, all of which generated considerably smaller datasets (14, 17, 18).

**Fig 2.**
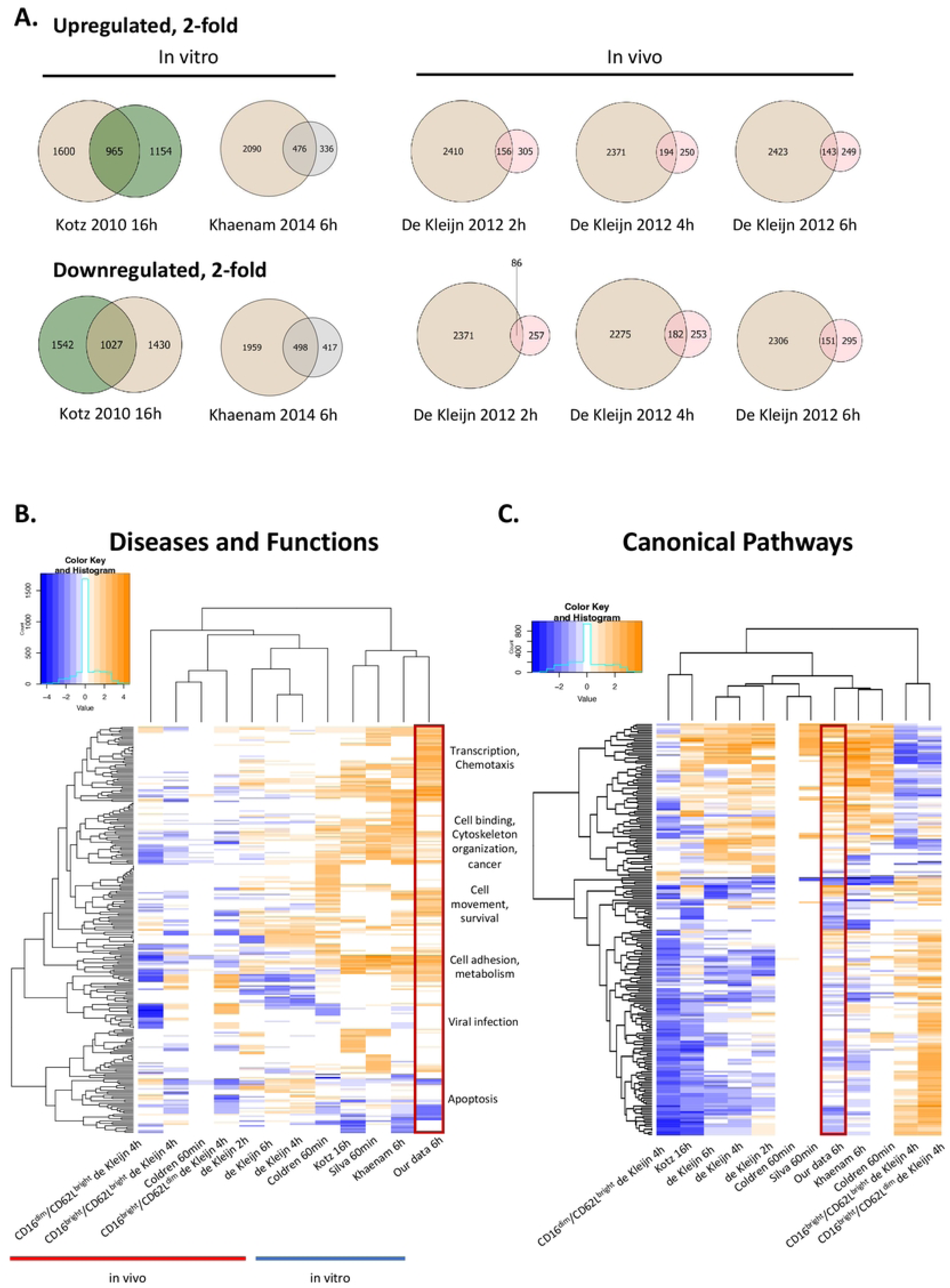
Meta-analysis of our data and other previously published LPS-treated neutrophil expression profiles. **A.** Venn diagrams illustrating partial overlap in genes regulated 2-fold in our dataset and other in vitro and in vivo datasets from LPS-stimulated neutrophils. Tan colour represents our 6h data and green, pink or grey colours represent previously published datasets, as indicated in the figure. Enriched diseases and biological functions **(B)** and canonical pathways **(C)** of genes in our dataset compared to other datasets from LPS-stimulated neutrophils (-log p value ≤ 1.3). Categories with a predicted activation state (positive Z-score) are labeled in orange while those with a predicted inhibition state (negative Z-score) are indicated in blue. Hierarchical clustering is shown to reveal similarly enriched datasets to our data. Our data is enclosed in the red rectangle.

**Fig 3.**
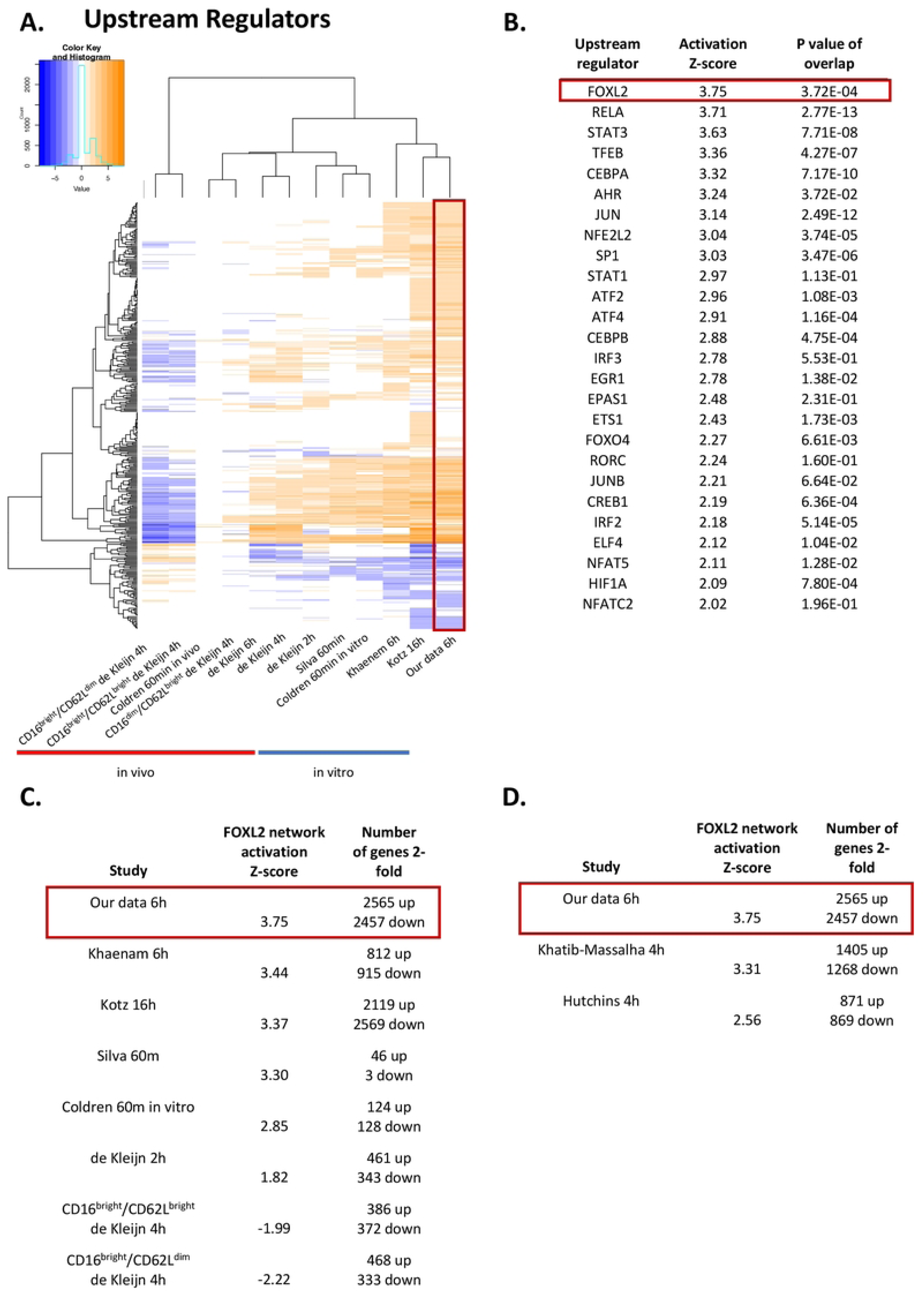
The FOXL2 transcriptional network is the most activated among transcription factor upstream regulators. **A.** Enriched upstream regulators of genes in our dataset compared to other datasets from LPS-challenged neutrophils. **B.** List of top activated upstream regulators that are transcription factors found in our data. FOXL2 is the most robustly enhanced transcriptional network, as indicated by the highest value for activation Z-score. The FOXL2 transcriptional network is conserved across several of the in vitro previously published human **(C)** and mouse **(D)** LPS-stimulated neutrophil studies. The number of 2-fold regulated genes for each dataset is shown.

We are interested in determining the transcription networks induced downstream of LPS stimulation. In our study, the most strongly enriched network was originally identified as being driven by the forkhead box transcription factor FOXL2 (Fig 3B) (35). Notably, the Z-score for the network is greater than those driven by RELA and STAT3, previously identified as regulators of LPS-induced inflammation in neutrophils (9, 10). Moreover, the FOXL2 network is conserved among other human and mouse *in vitro* LPS-stimulated neutrophil datasets (Figs. 3C, D). In contrast, it is not upregulated in two neutrophil subsets derived from cells exposed to LPS *in vivo* (Fig 3C). However, the vast majority of upstream regulators appear downregulated in these datasets (Fig 3A). In our data, 20 of the 22 genes in the network were induced, consistent with activation of a transactivator (Fig 4A). The network is based on the identification of transcription targets of FOXL2 in a human ovarian granulosa-like tumour (KGN) cell line (35). Nonetheless, numerous genes in the network encode proteins implicated in neutrophil function, and among those are chemokines, such as *CCL20* and *CXCL2* (36, 37) as well as stress and inflammation-associated transcription factors, *MAFF* and *ATF3* (38–42) (Fig 4A). Other genes in the network encode regulators of neutrophil survival, such as *BCL2A1* and *IER3*, respectively (43, 44) (Fig 4A). Genes implicated in other aspects of neutrophil function are also enriched. These include *ICAM1*, which promotes neutrophil adhesion and transcellular migration (45); *SOD2*, a mitochondrial enzyme capable of inducing neutrophil burst, leading to intracellular killing of pathogens (46); and *NR4A3*, an orphan nuclear receptor that boosts neutrophil numbers and survival (47) (Fig 4A). We validated increased expression of several of these genes by RT/qPCR analysis in neutrophils challenged with LPS (Fig 4B). Collectively, these findings suggest that the transcription network influences several aspects of neutrophil function and survival.

**Fig 4.**
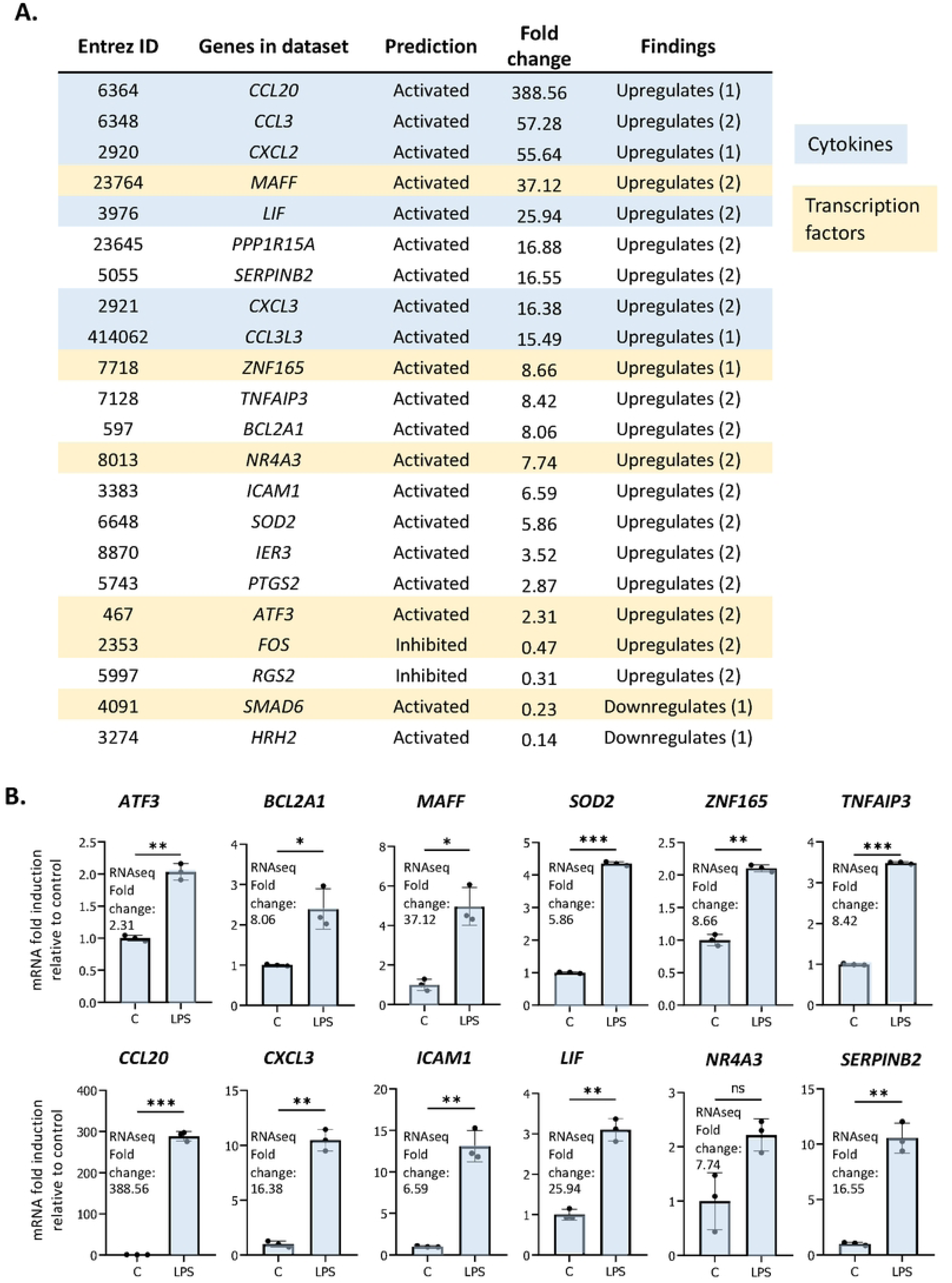
The FOXL2 network is enriched in genes encoding inflammation-associated transcription factors, cytokines and regulators of neutrophil function. **A.** All genes included in the network with corresponding Entrez ID, activated or inhibited prediction, fold change gene expression relative to control and whether the gene is up or downregulated based on previous findings in the literature. Genes that are cytokines are highlighted in blue, and genes that represent transcription factors are in beige. **B.** RT/qPCR analysis of LPS-regulated expression of genes within the FOXL2 network in human neutrophils. Graphics representative of 2 or 3 biological replicates. Graphics are mean ± SD from 3 technical replicates from a representative sample and paired, two-tailed t-test (Student’s t-test) was used (*P ≤ 0.05, **P ≤ 0.01, ***P ≤ 0.001 and ns ≥0.05).

### Induction of FOXP1 and FOXP4 expression in LPS-exposed neutrophils

Although *FOXL2* is not expressed in our dataset, forkhead box transcription factors *FOXP1* and *FOXP4* are expressed and induced by LPS in primary human neutrophils (22) (Fig 5A). Given that FOX transcription factors have highly conserved DNA binding domains, we hypothesized that FOXP1 and/or FOXP4, rather than FOXL2, drive expression of the network of genes in neutrophils (20). Previous studies revealed the importance of FOXP1 and FOXP4 in lymphocyte development and effector cytokine production, respectively (24–27), and FOXP1 regulates monocyte differentiation (23). We compared FOXP1 motifs from the JASPAR 2020 database with those of FOXL2, as determined by Carles et al (48) (Fig 5B), which revealed that FOXP1 and FOXL2 recognize essentially identical sequence “TGTAAACA” motifs, with the exception of the variable 5’ end of the sequence (Fig 5B). (Note that FOXP4 motifs are not identified in the JASPAR database). We confirmed increased gene and protein expression of the two proteins in LPS-stimulated neutrophils by RT/qPCR and Western blot analyses (Figs. 5C, D). Subsequently, FOXP1/4 ChIPseq peaks were identified within 14 of the 20 genes in the FOXL2 network using published ChIPseq studies and datasets from the ENCODE consortium (49, 50) (Fig 5E, Table S2). To find sequence motifs enriched in enhancers, we extracted their sequence from the hg19 or hg38 genome and used this as input for the Transcription factor Affinity Prediction (TRAP) web tool (http://trap.molgen.mpg.de/cgi-bin/home.cgi) (see Methods and Materials for details) (S2 File). The frequency of the FOXP1/P4 peaks in the neutrophil network genes was remarkable given that the ChIPseq studies were performed in the DLD1 colon and HepG2 liver cancer cell lines, as well as in embryonic stem cells (49, 50) (Table S2). FOX protein consensus GTAAACA or near-consensus motifs were present at multiple binding sites in 8 of the 14 network genes containing ChIPseq peaks (see Fig 5E). By comparison, the van Boxtel et al study (50) identified TGTTTAC, the reverse complement sequence of GTAAACA, motifs in the vicinity of 30% of FOXP1 ChIPseq peaks. Notably, genes that contain the FOX protein motifs included inflammation-induced transcription factors MAFF, ATF3, and ZNF165 and those encoding several cytokines (Fig 5E).

**Figure 5.**
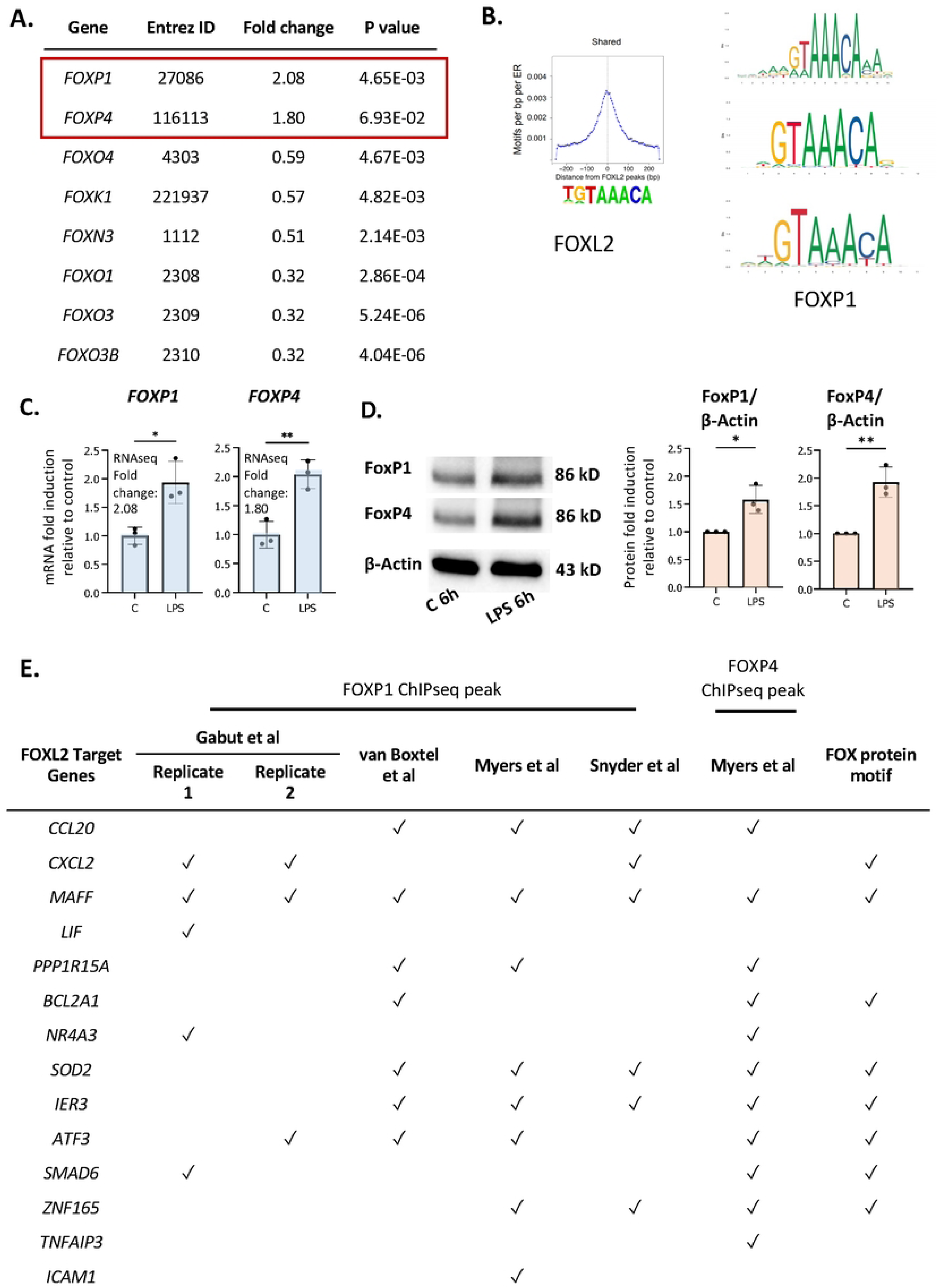
FOXP1 and FOXP4 binding sites are present in regulatory regions of network genes. **A.** Up- and downregulated genes by LPS encoding FOX transcription factors in our RNAseq data. **B.** Conserved motifs (sequence logos) of FOXL2 (left, retrieved from Carles et al (48)) and FOXP1 (right, retrieved from the JASPAR 2020 database). **C.** RT/qPCR analysis of LPS-regulated expression of genes encoding the FOX transcription factors, FOXP1 and FOXP4. Data representative of 2 or 3 biological samples. Graphics are mean ± SD from 3 technical replicates from a representative sample and paired, two-tailed t-test (Student’s t-test) was used (*P ≤ 0.05 and **P ≤ 0.01). **D.** Western analysis of FOXP1 and FOXP4 protein expression in vehicle- or LPS-treated neutrophils, as indicated. Data representative of 3 biological replicates. Quantification shown on right. Graphics are mean ± SD from 3 biological replicates and unpaired, two-tailed t-test was used (*P ≤ 0.05 and **P ≤ 0.01). **E.** Presence (marked by a tick) or absence of FOXP1 and/or FOXP4 ChIPseq peak(s) and FOX protein motifs within the network genes in different FOXP1 or FOXP4 ChIPseq datasets performed in various cell types.

To further probe whether the FOXP1/P4 ChIP binding sites may be active in neutrophils, we identified regions of histone marks in neutrophils corresponding to enhancers (regions of H3K4 trimethylation and H3K27 acetylation) from datasets from the ENCODE consortium (Figs. 6A, S6, Table S2). Notably, the binding sites containing motifs do not overlap areas of H3K27 or H3K9 trimethylation (Figs. 6A, S6), which are histone marks indicative of transcriptionally repressed regions. Furthermore, we found that the promoter-proximal FOXP1/P4 binding sites overlap with RNA polymerase II binding profiles in neutrophils, as illustrated in the UCSC browser image for the *ZNF165* locus (Fig S6). Two clusters of FOXP1/4 binding sites are present in the vicinity of the *MAFF* gene; one at approximately 14 kb upstream from the transcription start site (TSS), and another in an intronic region of the gene (∼700 bp from the TSS) (Figs. 6A, S6). Furthermore, a promoter-proximal FOXP1 binding site at 221 bp downstream from the TSS containing a FOX protein motif is present in the *ZNF165* gene (Fig S6). Taken together, these results strongly suggest that, in LPS-treated cells, putative FOXP1 and FOXP4 binding sites are present in regions corresponding to neutrophil enhancers.

**Figure 6.**
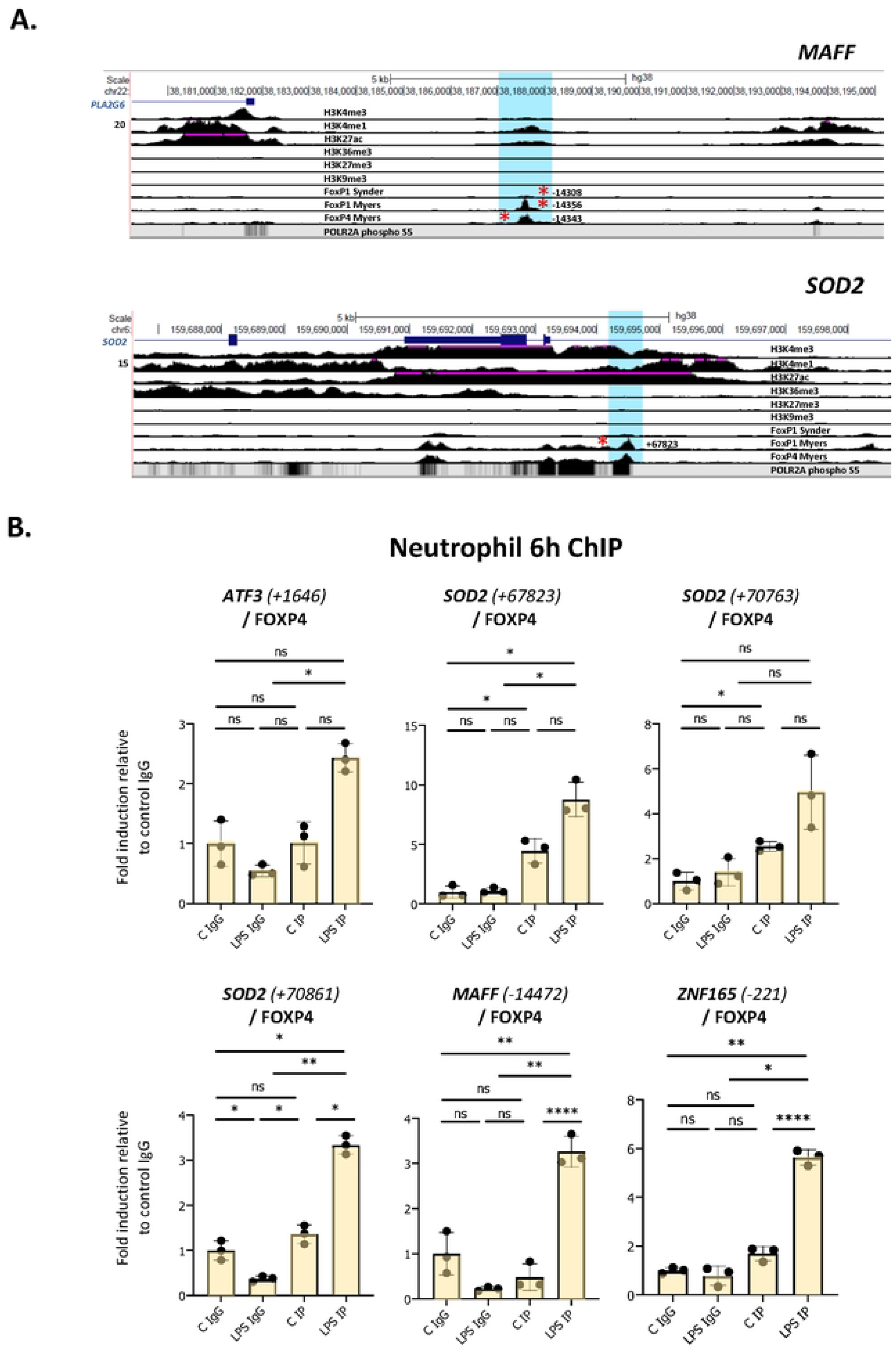
Enhanced binding of FOXP4 to network gene motifs as assessed by ChIP assay in LPS-challenged neutrophils. **A.** UCSC browser images showing FOXP1 and FOXP4 ChIPseq tracks at MAFF and SOD2 loci. The areas surrounding the FOXP1/FOXP4 binding sites up and downstream (regions highlighted in blue) are shown. Red asterisks represent ChIPseq peaks that contain FOX protein motifs. Binding sites correspond to regions of enhancer function in neutrophils. **B.** Analysis of the association of FOXP4 with up- and downstream regulatory regions of ATF3, SOD2, MAFF and ZNF165 by ChIP assay in neutrophils treated with or without LPS for 6h. Data representative of at least 3 biological replicates. Graphics are mean ± SD from 3 technical replicates from a representative sample. *P ≤ 0.05, **P ≤ 0.01, ***P ≤ 0.001, ****P ≤ 0.0001 and ns ≥0.05 as assessed by one-way ANOVAs followed by Tukey’s post hoc test for multiple comparisons. ChIP values are normalized to input for each condition and expressed as a fold relative to non-specific IgG control.

To determine whether FOXP1 and/or FOXP4 interact with these sites in neutrophils, we performed ChIP assays in vehicle- and LPS-treated cells. These experiments were performed in multiple isolates of primary human neutrophils. We observed elevated binding of FOXP4 induced by LPS to multiple sites in primary cells (Fig 6B, Fig S7A). Enhanced FOXP1 binding was much more difficult to detect. While we did observe elevated binding in the presence of LPS to multiple sites in one neutrophil isolate (Fig S7B), these findings were not reproducible in other primary human cells (data not shown).

### Binding of forkhead box proteins to multiple network regulatory sites in transfected HEK293 cells and activation of network genes

Neutrophils are not stable in tissue culture and problematic to manipulate genetically. Therefore, to further investigate the potential roles of forkhead box proteins in binding to enhancer regions described above and in the regulation of network target genes, we used HEK293 cells, a heterogeneous cell system that does not express either of the two proteins (Fig 7A). FOXP1 and FOXP4 expression was observed in cells transiently transfected with corresponding expression vectors (Fig 7A). ChIP assays were performed to analyse binding of the two proteins to multiple binding sites identified above. FOXP4 binding was observed to all sites (Fig 7B). Remarkably, robust binding of FOXP1 by ChIP was also observed to all sites (Fig S7C), with the strongest signals at sites adjacent to the *ATF3* and *SOD2* genes. Importantly, we also observed a general increase in mRNA expression of multiple network genes in transfected HEK293 cells (Fig 7C). Taken together, these data show that expression of network genes is induced under conditions of elevated forkhead box transcription factor expression.

**Figure 7.**
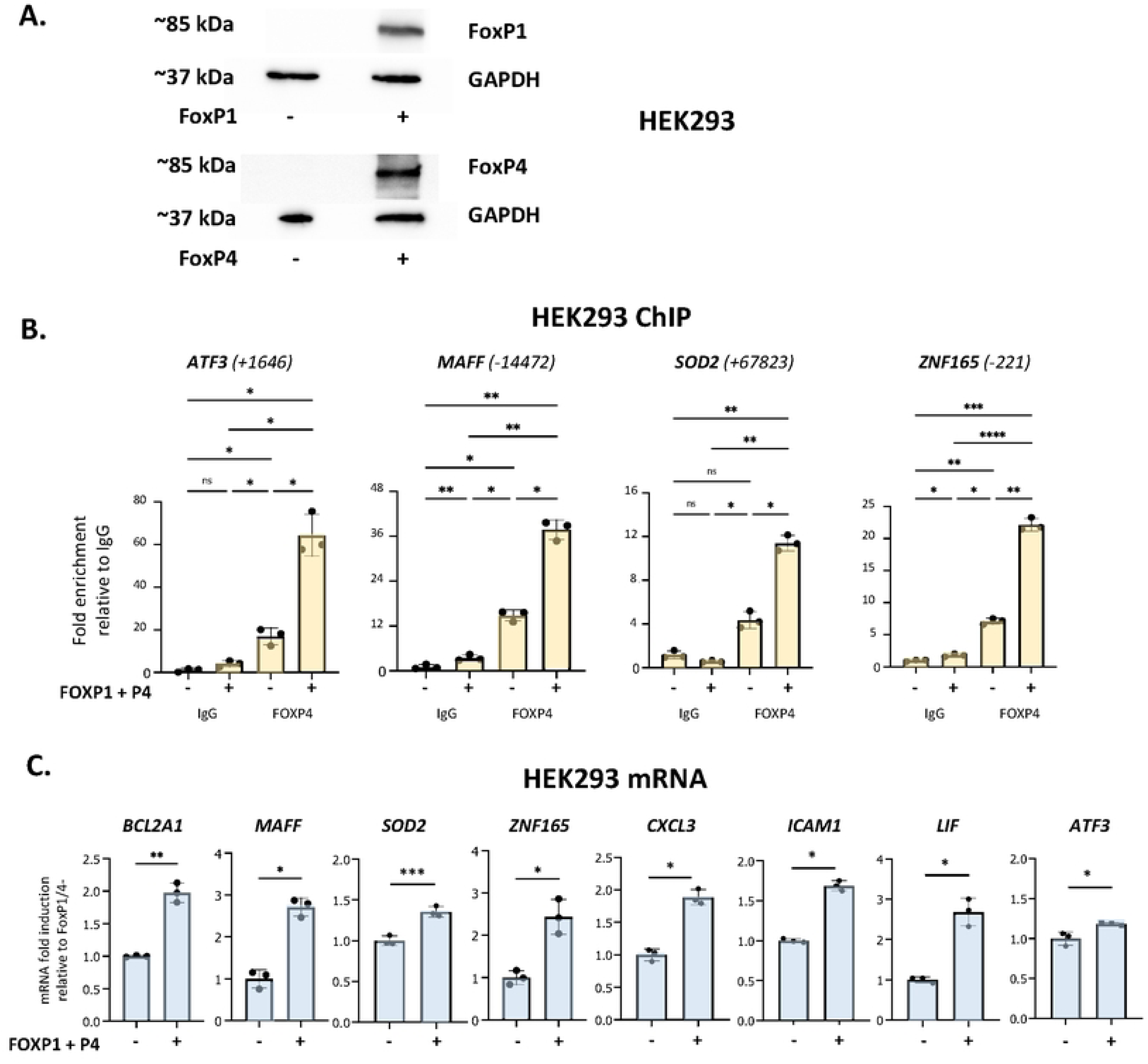
Analysis of forkhead box protein DNA-binding and gene regulation in HEK293 cells. **A.** Western analysis of FOXP1 and FOXP4 protein expression in HEK293 cells transfected with or without FOXP1 and FOXP4 expression vectors, as indicated. **B.** Analysis of the association of FOXP4 with up- and downstream regulatory regions of ATF3, MAFF, SOD2 and ZNF165 by ChIP assay in HEK293 cells transfected with and without FOXP1 and FOXP4 expression vectors. Data representative of 2 or 3 biological replicates. Graphics are mean ± SD from 3 technical replicates from a representative sample and paired one-way ANOVAs followed by Tukey’s post hoc test for multiple comparisons were used (*P ≤ 0.05, **P ≤ 0.01, ***P ≤ 0.001, ****P ≤ 0.0001 and ns ≥0.05). ChIP values are normalized to input for each condition and expressed as a fold relative to non-specific IgG control. **C.** RT/qPCR analysis of FOXP1/4 network genes in HEK293 transfected with and without FOXP1 and FOXP4 expression vectors. Data representative of 2 or 3 biological replicates. Graphics are mean ± SD from 3 technical replicates from a representative sample and paired, two-tailed t-test (Student’s t-test) was used (*P ≤ 0.05, **P ≤ 0.01, ***P ≤ 0.001, ****P ≤ 0.0001 and ns ≥0.05).

We next investigated whether induction of the forkhead box protein network is observed in neutrophils stimulated with inflammatory agents other than LPS. The GEO repository was used to search for expression profiles of human neutrophils challenged with inflammatory signals, which generated 4 microarray datasets (see Table S3 for details) (16, 47, 51, 52). They included neutrophils challenged with the protein kinase A agonist pair 8-AHA-cAMP and N6-MB-cAMP (N6/8-AHA) and cytokines GM-CSF and interferon-gamma (IFN-γ) over time periods ranging from 3-24h (Table S3) (16, 47). Analysis of these datasets revealed that the FOX protein network was enriched in all profiles with a high Z-score (Fig 8).

**Figure 8.**
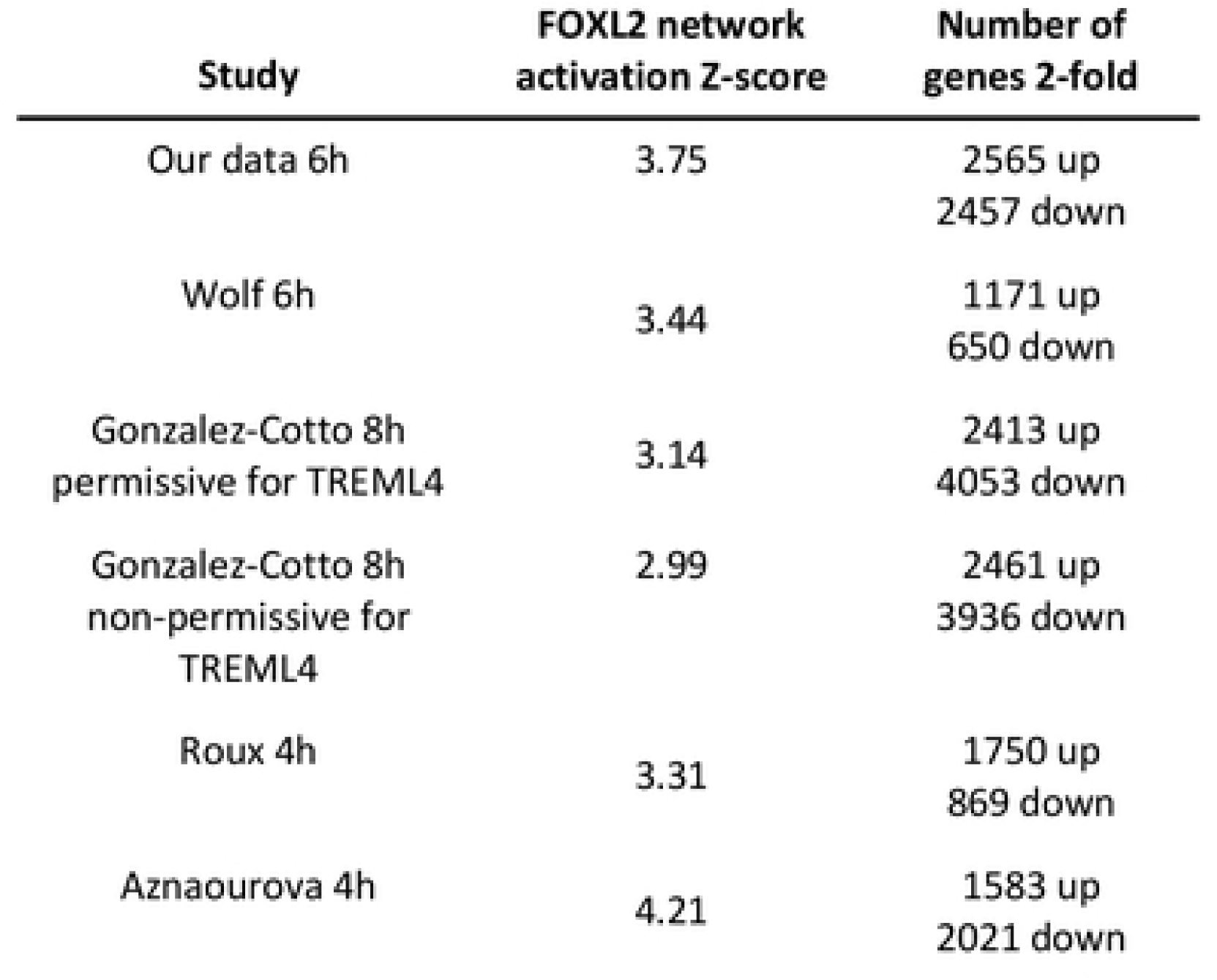
FOX protein network is induced by LPS in neutrophils in response to other inflammatory signals. Lists of other datasets of human neutrophils stimulated with various human inflammatory signals. Their associated FOX network activation Z-scores and number of 2-fold regulated genes are shown as well.

## Discussion

To date, analysis of transcriptional changes occurring in neutrophils challenged with LPS has revealed activation of NF-κB and STAT3 transcription factors (9, 10) as well as increased pro-inflammatory cytokine expression (11–19). Additional research into such changes provides insight into novel signaling pathways and transcriptional responses driving in neutrophil immune responses under (patho)physiological conditions. In this study, we performed the first large scale RNAseq profile to probe the human neutrophil transcriptomic responses to LPS and other inflammatory agents. Bioinformatic analysis of this data showed that treatment with LPS robustly induced a transcriptional network, initially identified as being regulated by the forkhead transcription factor FOXL2 in other cell types. However, the network is relevant to neutrophil function as it is highly enriched in genes encoding proinflammatory cytokines and transcription factors implicated in inflammatory and stress responses. Importantly, the network includes several genes encoding transcription factors such as ATF3 and MAFF, which have been implicated in cellular responses to stress and inflammatory signals (38–42), as well as that encoding the orphan nuclear receptor NR4A3, whose expression controls neutrophil numbers and survival (47). The network thus controls a cascade of transcriptional events in response to LPS.

While FOXL2 was not expressed in our RNAseq dataset, forkhead box transcription factors FOXP1 and FOXP4 were expressed and induced by exposure to LPS. Several lines of evidence indicate that FOXP4 can functionally replace FOXL2 in neutrophils. These include the fact that members of the forkhead box transcription family recognize essentially identical DNA motifs; all FOX transcription factors have a conserved DNA binding domain and bind to essentially identical GTAAACA motifs (20, 48). Moreover, FOXP1/4 binding sites are present in network genes identified in ChIPseq studies performed in heterogeneous cell types, but located in regions that correspond to transcriptional enhancers in neutrophils. Further bioinformatics analysis revealed that the genes in the network were also regulated in neutrophils exposed to other inflammatory signals.

We performed ChIP assays on multiple isolates of primary human neutrophils and observed enhanced LPS-dependent binding of FOXP4 to several motifs adjacent to network genes. While we did find some evidence for LPS-induced FOXP1 binding in one neutrophil isolate, this finding was not generally reproducible. Studies in transiently transfected HEK293 cells showed that both FOXP1 and FOXP4 can recognize multiple motifs in network genes. The binding of FOXP1 to these motifs in HEK293 cells may be a function of its elevated expression under conditions of transient transfection. However, collectively our data suggest that we cannot rule out the possibility that FOXP1 may contribute to network gene transcription if sufficiently induced in neutrophils. Moreover, the data in HEK293 cells showed that forkhead box transcription factor expression can induce transcription of multiple network genes.

Meta-analysis of our RNAseq data and other available comparable expression profiles revealed that canonical pathways, diseases and functions, and upstream regulators identified in other *in vitro* profiles clustered closer to our data than those derived from neutrophils exposed to LPS *in vivo*. A high activation Z-score was observed for the FOXL2 transcriptional network in all *in vitro* datasets, which is remarkable given that the meta-analysis included expression profiles performed using different platforms, and with neutrophils exposed to LPS for varying times and at different concentrations. However, the network displayed a negative Z-score for two neutrophil subpopulations acquired from cells stimulated with LPS *in vivo* (18). This may be due partly to cell sorting by FACS performed on the cells treated *in vivo*, which may perturb gene expression profiles (54). The discrepancies may also be explained by the method in which neutrophils were treated with LPS. *In vitro* neutrophil studies consist of isolating cells from human whole blood (14–16, 19). In contrast, in *in vivo* experiments, a human endotoxemia model (intravenous administration of an endotoxin solution) was employed (14, 17, 18). There is a limitation to this model as it produces a low-grade acute systemic inflammatory state and does not completely recapitulate chronic inflammation present in inflammatory diseases (55). The LPS dose *in vivo* required to stimulate, for example, sepsis (an acute inflammatory response to infection in which neutrophil function is dysregulated) is unsafe and ethically unacceptable in human studies (56). Moreover, the dynamic alterations in cytokine levels noted after LPS injection are substantially different from the more sustained levels observed in critically ill patients (56) and *ex vivo* experiments (57). Finally, the single-exposure human endotoxemia model cannot capture sustained innate immune activation that may occur with chronic exposure to inflammatory agents (55).

In addition to LPS, the network identified here may be regulated by other upstream regulators or stimuli. We found that the network was induced in response to other neutrophil inflammatory agents, such as the protein kinase A agonist pair N6/8-AHA and cytokines GM-CSF and IFN-γ. In addition, bioinformatic analysis of gene expression in neutrophils treated with two different strains of bacteria revealed a high activation Z-score for the network. Germline deletion of FOXP1 or FOXP4 leads to embryonic lethality in mice due to cardiac defects (22, 61). Other studies have investigated the role of the two proteins in T and B lymphocytes using conditionally targeted strains in rodents (24–27). Feng et al crossed floxed FOXP1 mice with Cd4^Cre^ transgenic mice (24, 25). These conditional knockout studies found that FOXP4 and FOXP1 are implicated in the regulation of B and T cell development and cytokine production. Additionally, FOXP1 is expressed in untreated and retinoic acid-induced differentiated HL-60 cells; its expression was inhibited in phorbol ester-induced HL60 monocytic differentiation but not in retinoic acid-induced granulocytic differentiation (23). Collectively, therefore, our findings and evidence from their roles in regulation of other immune cell types suggest that FOXP1 and/or 4 may play key roles in neutrophil biological functions. Conditional knockouts of FOXP1 and/or FOXP4 would be a future avenue of study to better understand the roles of the two proteins in neutrophil differentiation and function. In conclusion, our study identifies a novel transcription network induced by forkhead box transcription factors in neutrophils exposed to inflammatory stimuli. Our findings lay the foundation for additional investigations into the roles of FOXP4 and FOXP1 in neutrophil biology. Further research using appropriate animal models are required to assess their function and the role of the network in neutrophil-associated inflammatory diseases, such as rheumatoid arthritis and cancer.

## Materials and Methods

### Human neutrophil isolation and treatment

Whole blood from consenting healthy donors was provided by Dr. Jack Antel (McGill) through C-Big under McGill University Health Centre REB ethics #2021-6588. Primary human neutrophils were isolated from blood by negative selection using the EasySep™ Direct Human Neutrophil Isolation Kit (STEMCELL) according to the manufacturer’s instructions. Purity of neutrophils was determined by flow cytometry by quantifying markers of various cell populations found in blood, notably CD45 (hematopoietic cells with the exception of erythrocytes and platelets), CD16 (natural killer cells, neutrophils and macrophages) and CD66b (granulocytes). Cells were counted by an automatic cell counter (Bio-Rad) and adjusted to a concentration in between 5 × 10^5^ and 1 × 10^6^ cells/ml. Neutrophils were resuspended in tissue culture medium, which consisted of RPMI 1640, 1X with L-glutamine, sodium pyruvate & 25mM HEPES (350-006-CL, Wisent) supplemented with 10% fetal bovine serum and penicillin/streptomycin (0503, ScienCell). Cells were subsequently treated with 1µg/mL LPS (L3012-5MG, Sigma-Aldrich) or vehicle (dimethyl sulfoxide) for 6 hours. By annexin V/propidium iodide staining, we showed that after 6h neutrophils were mostly viable (Fig S8).

### Data collection

We mined the PubMed database for microarray and RNAseq expression profiling. We used the following key words and their combinations: “Neutrophil, LPS, lipopolysaccharide, microarray, RNAseq, gene expression dataset”. Furthermore, the Gene Expression Omnibus (GEO) public repository was used to search for studies using the keywords “lipopolysaccharide AND neutrophil.” Gene expression studies investigating the effects of lipopolysaccharide on neutrophils, for which the raw data were available, were included. We extracted the following information from each identified study: publication reference, GEO accession number, dose and time of LPS treatment, platform, and number of biological replicates (Table S1). The analysis excluded studies in LPS knockout mice.

### RNA sequencing

Total RNA from LPS- and vehicle-treated neutrophils was extracted using FavorPrep Blood/Cultured Cell Total RNA Mini Kit (FABRK 001, Favorgen) as per the manufacturer’s instructions. Biological replicates were generated from 3 independent neutrophil isolates. Only RNA samples with OD 260/280 ratio greater than 1.7 and a RNA integrity number (RIN) > 7 were kept for downstream analysis. These samples were submitted to McGill University and Genome Quebec Innovation Centre for paired-end sequencing at 50M reads by Illumina NovaSeq 6000 S2 PE100. Quality control including read trimming and filtering out low quality reads, mapping to GRCh38 genome assembly, summarization by gene, normalization and differential gene exression using the R packages EdgeR and DESeq2 was performed at McGill University and Genome Quebec Innovation Centre. Running the RNAseq quality control pipeline, it was determined that 3 and 2 replicates for control and LPS-treated conditions, respectively, were of high enough quality for downstream analysis. We obtained read counts and assessed differential gene expression using EdgeR and LIMMA R packages, respectively. Differentially-expressed genes were 2-fold regulated and p value ≤ 0.05 was used as the threshold for significance. Principal component analysis (PCA) was performed using the built-in R function, prcomp, and visualized with the ggplot package (Fig S3).

### Flow cytometry

Adherent neutrophils were detached by gently pipetting the tissue culture dishes up and down. Adherent and suspension cells were centrifuged at 500 rcf for 10 min, and washed twice with ice-cold PBS. The supernatant was subsequently removed. We resuspended the cells in FACS buffer (0.5-1% BSA in PBS) at a concentration of 1 × 10^6^ cells/ml and blocked the cells with human FcR binding inhibitor (14-9161-73, eBioscience). To determine neutrophil purity, 2 µg of anti-human PerCP/Cy5.5-CD66b (305107, BioLegend), PE-CD16 (302007, BioLegend), and PE/Cy7-CD45 (103113, BioLegend) antibodies were added and incubated for 30 min at room temperature in the dark. Viability was assessed using the Vybrant Apoptosis Assay kit (V13242, Molecular Probes). Cells were washed and either cross-linked in 2% paraformaldehyde or immediately analyzed by flow cytometry for purity and viability experiments, respectively. A BD-LSRFortessa analyzer was employed for flow cytometry acquisition and at least 10 000 cells/sample were monitored. The FlowJo software (TreeStar Inc.) was subsequently used for data analysis.

### Meta-analysis of gene expression profiles

Analyses of gene expression profiles was performed using the R statistical package. The oligo package was employed to receive signal intensities from CEL files of Affymetrix microarrays; data was normalized and summarized using the Robust Multi-Array Average method, a component of the oligo package. Using Illumina’s BeadStudio, Illumina raw input was summarized. The LIMMA package in R was employed to normalize Illumina, Agilent, Nimblegen, and custom arrays. RNAseq raw files (SRA) were downloaded and transformed to reads in fastq format by the fastq-dump function in the sratoolkit suite from NCBI. A genome index was constructed derived from human/mouse genomes, supplied by Guillimin, Calcul Quebec high-performance computing cluster at McGill University, using the Rsubread package. The latter was also used to align RNAseq reads to retrieve read counts based on the “seed-and-vote” paradigm. The ensuing data was arranged for downstream analysis using the package EdgeR. For assessing differential gene expression for microarray and RNAseq data, the LIMMA package was employed. Differentially-expressed genes were 2-fold regulated and p value ≤ 0.05 was used as the threshold for significance. The biomaRt package, an interface to the BioMart databases at Ensembl, was used for annotation, including human orthologs for mouse genes.

### Quality control criteria

Each RNAseq dataset had similar total number of RNAseq reads sequenced for each condition and total GC content. The Phred offset quality score was greater than 30 for all RNAseq datasets and the minimum fragment size for alignment was set to 50. These settings yielded 70% to 85% of reads aligning to a gene using the RSubread and EdgeR packages in R. We filtered out genes with low read counts and included genes with counts of 10 or more for at least 1 treatment group (in all replicates) for downstream analysis. To validate similarity between replicates in each treatment group, we performed a principal components analysis (Fig S3). For Illumina microarray platforms, the probe summary files, which comprised the control probes, were exported from Illumina’s GenomeStudio without background correction or normalization. Raw files from the remaining microarray datasets were analyzed in R using the GEOquery, oligo and LIMMA packages. To determine potential outliers in microarray datasets, we visualized signal distribution of the raw and normalized data using box plots (Fig S4). Samples in all datasets seemed comparable following background correction and normalization.

### Bioinformatics analysis

Venn diagrams were generated with the VennDiagram package in R. Enriched canonical pathways, upstream regulators, biological diseases and functions as well as the FOXP1/4 upstream transcriptional network were identified using QIAGEN’s Ingenuity Pathway Analysis software (IPA®, QIAGEN Redwood City, www.qiagen.com/ingenuity). Heatmaps with hierarchical clustering were constructed using the heatmap.2 package in R. Peaks from FOXP1/4 ChIPseq studies and datasets from the ENCODE consortium were aligned with the human genome (build hg19 or hg39) using the UCSC Genome Browser (http://genome.ucsc.edu/cgi-bin/hgGateway). FOXP1 or FOXP4 binding sites were considered if they were within 25kb of a gene and in regions of active (either H3K4 trimethylation and H3K27 acetylation), but not repressive (H3K27 nor H3K9 trimethylation) histone marks in neutrophils using the statistical analyses provided in each publication. To find sequence motifs enriched in enhancers in human cancer cell lines and embryonic stem cells, we extracted their sequence from the hg19 or hg38 genome and used this as input for the Transcription factor Affinity Prediction (TRAP) web tool (http://trap.molgen.mpg.de/cgi-bin/home.cgi) using JASPAR vertebrates as the comparison library, human promoters as the control, and Benjamini-Hochberg as the correction (62). We used a p value threshold of 0.05. This resulted in the enrichment of several consensus or near-consensus motifs (S2 File) for FOX transcription factors. Matrices used were FOXO3: MA0157.1; FOXD1: MA0031.1; FOXF2: MA0030.1, FOXA1: MA0148.1, and FOXC1: MA0032.1.

### Cell culture

HEK293 cells were obtained from the American Type Culture Collection (ATCC) and cultured in DMEM (319-005-CL, Wisent) supplemented with 10% fetal bovine serum and penicillin/streptomycin.

### RNA extraction, reverse transcription and qPCR

We performed RNA extraction with the FavorPrep™ Tissue Total RNA Mini Kit (FATRK 001, Favorgen) as per the manufacturer’s instructions. cDNA was acquired from 100 - 500 ng of RNA using 5× All-in-One RT Mastermix (G485, abm) and diluted 5 times. Quantitative polymerase chain reaction (qPCR) was performed with BrightGreen 2×qPCR MasterMix (MasterMix-LR-XL, abm) on a Roche Applied Science LightCycler 96 machine. We normalized the expression of genes was normalized to *18S* or *ZC2HC1C*. All primers are listed in Table S4.

### Western blotting and protein analysis

Primary human neutrophils and HEK293 cells were solubilized in Lysis Buffer (20 mM Tris, pH 8, 150 mM NaCl, 1% Triton X-100, 3,5 mM sodium dodecyl sulfate, 13 mM deoxycholic acid) and extracted proteins were separated on a 4% to 15% Tris/Glycine/sodium dodecyl sulfate gel (Bio Rad). For transfer and blotting, we used a standard protocol. FOXP1 (#ab16645, abcam, 1:500) and FOXP4 (#ab17726, abcam, 1:500) primary antibodies were purchased from Abcam. The antirabbit IgG HRP-linked secondary antibody was purchased from Cell Signaling Technology and used at recommended concentrations. Signals of protein bands were detected using Clarity ECL chemiluminescent substrates (Bio-Rad) and ChemiDoc Imaging System (Bio-Rad). We quantified changes in protein levels relative to control using Image Lab software after normalization to β-actin or GAPDH (Cell Signaling Technologies), as indicated. Western blot images are representative of three biological replicates.

### Chromatin immunoprecipitation assays

Cells were cross-linked with 1% formaldehyde for 15 min and were lysed with 500 µl lysis buffer (20mM Tris HCl pH 8, 1% SDS, 50mM NaCl) containing 1x protease inhibitor cocktail (with EDTA). Chromatin samples were sheared to a length of 300-500 bp via sonication. Following centrifugation, supernatants were collected and 2 µg of antibody (Abcam, same as in Western blotting) was added to chromatin to immunoprecipitate overnight. Dynabeads Protein G (10003D, Thermofischer) was added to antibody chromatin complexes for 2h. Next, protein G bead-chromatin complexes were washed twice with dilution buffer (20mM Tris HCl pH 7.6, 1% Triton X-100, 150mM NaCl) and once with washing buffer (PBS, 0.02% Tween-20, pH 7.4), as specified by manufacturer’s instructions. Proteinase K (P8107S, New England Biolabs) was added to immunoprecipiated and input chromatin; samples were then incubated at 45°C for 2h. Subsequently, chromatin was heated at 64°C for 4h for reversal of formaldehyde crosslinking. DNA fragments were purified using a PCR purification kit (FAGCK001-1, Favorgen) and were analyzed by qPCR. Antibodies used for ChIP are the same as for western blotting. Primer pairs used for ChIP assays are listed in Table S4.

### Statistics

Two-tailed t-test (Student’s t-test), performed using GraphPad software, was used to determine significance of results for 2 conditions. For 4 conditions, a one-way ANOVA followed by Tukey’s post hoc test for multiple comparisons was used using GraphPad. A p value of less than or equal to 0.05 was considered significant. To denote p values, symbols were used as follows: **P ≤ 0.05, **P ≤ 0.01, ***P ≤ 0.001, ***P ≤ 0.0001, and ns ≥0.05. Results from RT/qPCR, western blotting and ChIP analyses are representative of at least 3 biological replicates and one-way ANOVAs were used to determine significance. Paired tests were used for technical replicates of a representative sample and unpaired tests were used for biological replicates.

## Acknowledgments

We are grateful to Drs. Jack Antel, Luda Diatchenko, Jörg Fritz, Jo Stratton, Jelani Clarke and Catie Futhey as well as Nicholas Kieran and Maha Zidan for providing primary human neutrophils. We thank the Flow Cytometry and Cell Sorting Facility at McGill, as well as Julien Leconte and Camille Stegen, for assistance with the flow cytometry experiments; the facility’s infrastructure is supported by the Canada Foundation for Innovation (CFI).

## Author Contributions

Wrote the paper: AI, JHW. Conceived and designed the experiments: JHW, AI. Performed the experiments: AI, VD, CB. Methodology of ChIP assays and HEK293 transfections: RST, BM. Analyzed the data: AI. Methodology of bioinformatic analyses: AI, VD. Supervision: JHW.

## Supporting information

**S1 Fig.**
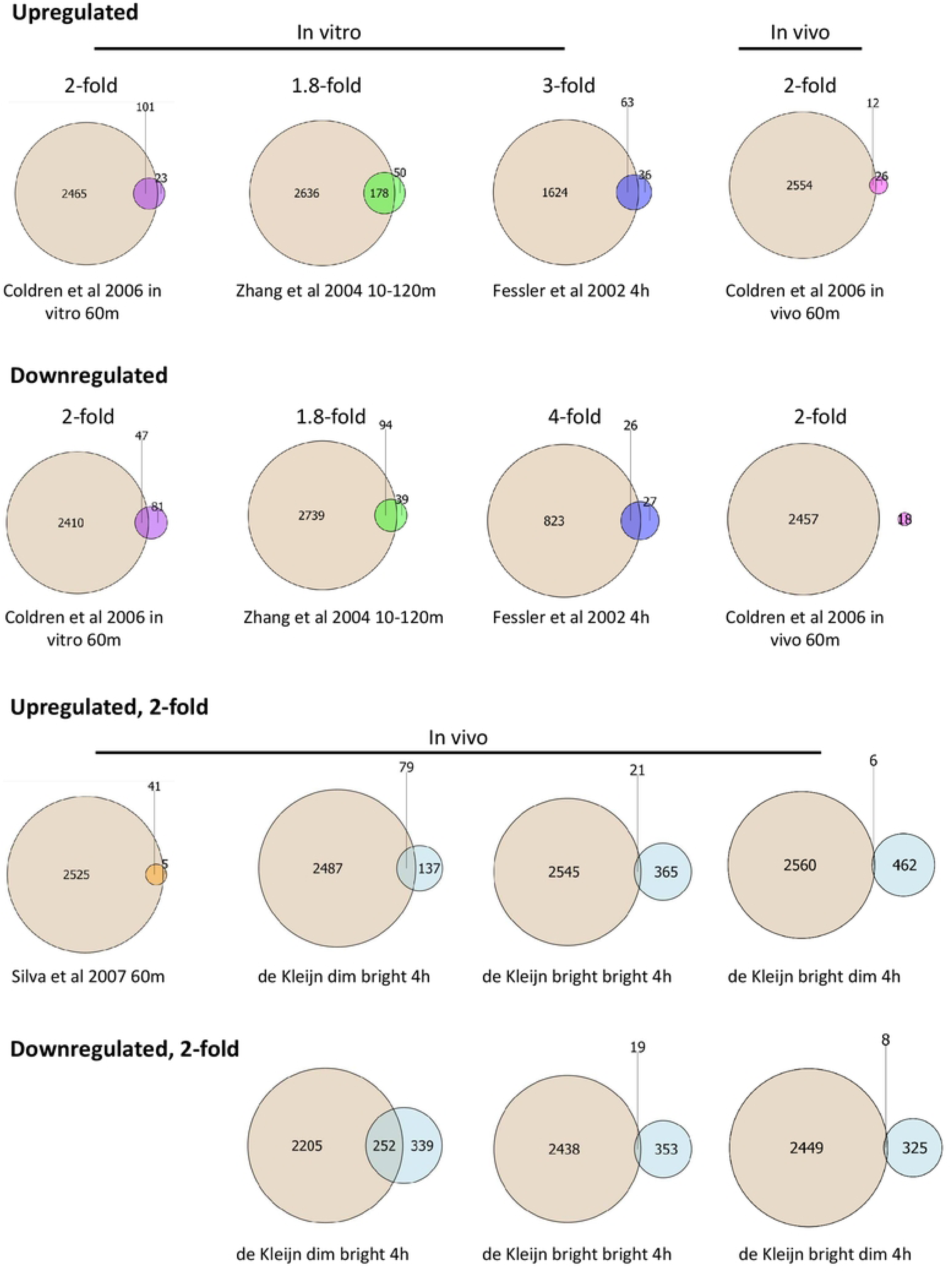
Venn diagrams illustrating partial overlap in genes regulated 2-fold in our dataset and other in vitro and in vivo datasets from LPS-stimulated neutrophils. Tan colour represents our 6h data and other colours represent previously published datasets, as indicated in the figure.

**S2 Fig.**
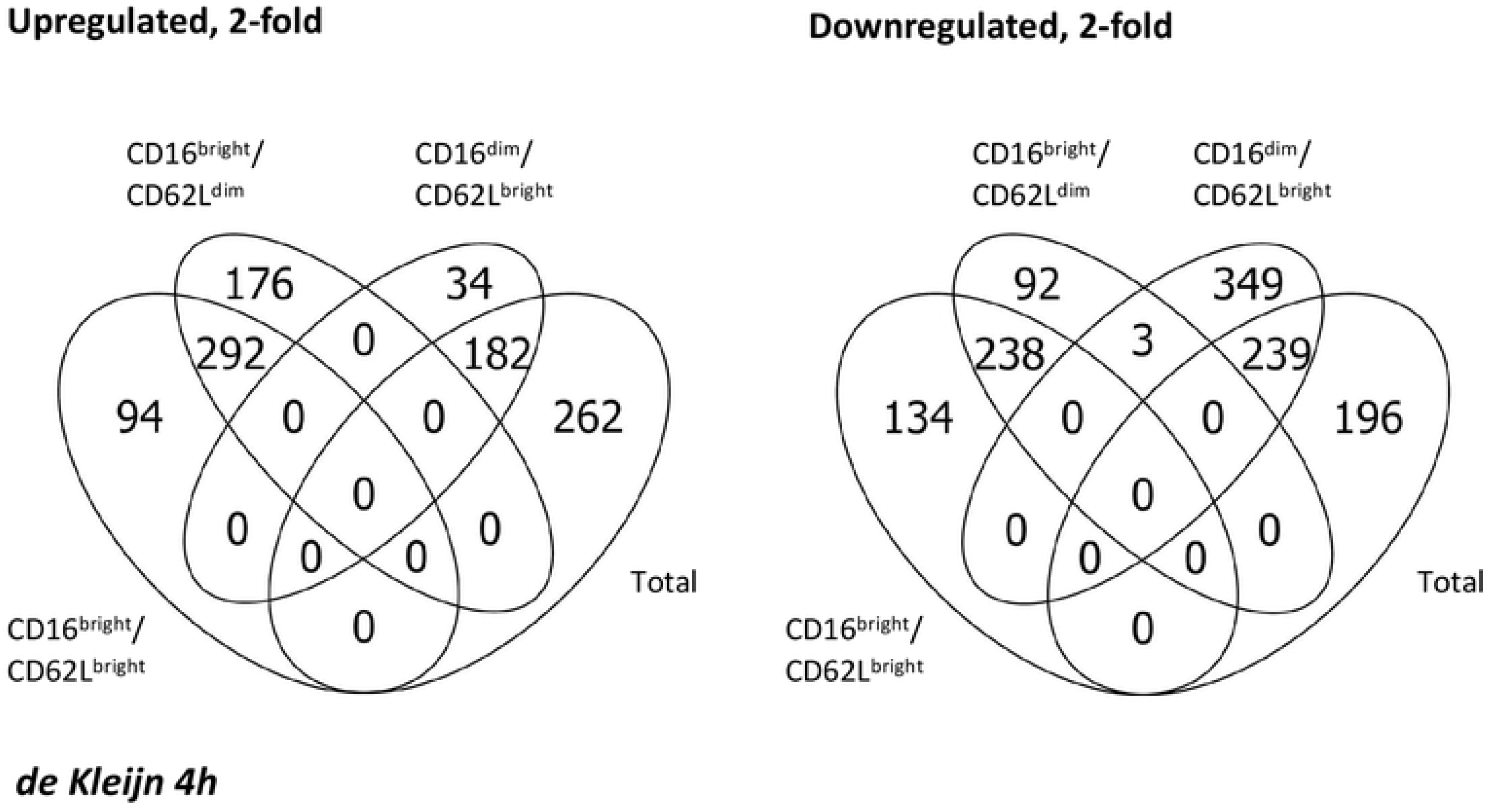
Venn diagrams of 3 sorted subpopulations and total neutrophils from 4h LPS-stimulated neutrophil datasets from de Kleijn et al 2013 and 2012, respectively. Little to no overlap is illustrated between the subpopulations and total cells, except for a partial overlap observed for total and CD16^dim^/CD62L^bright^ cells.

**S3 Fig.**
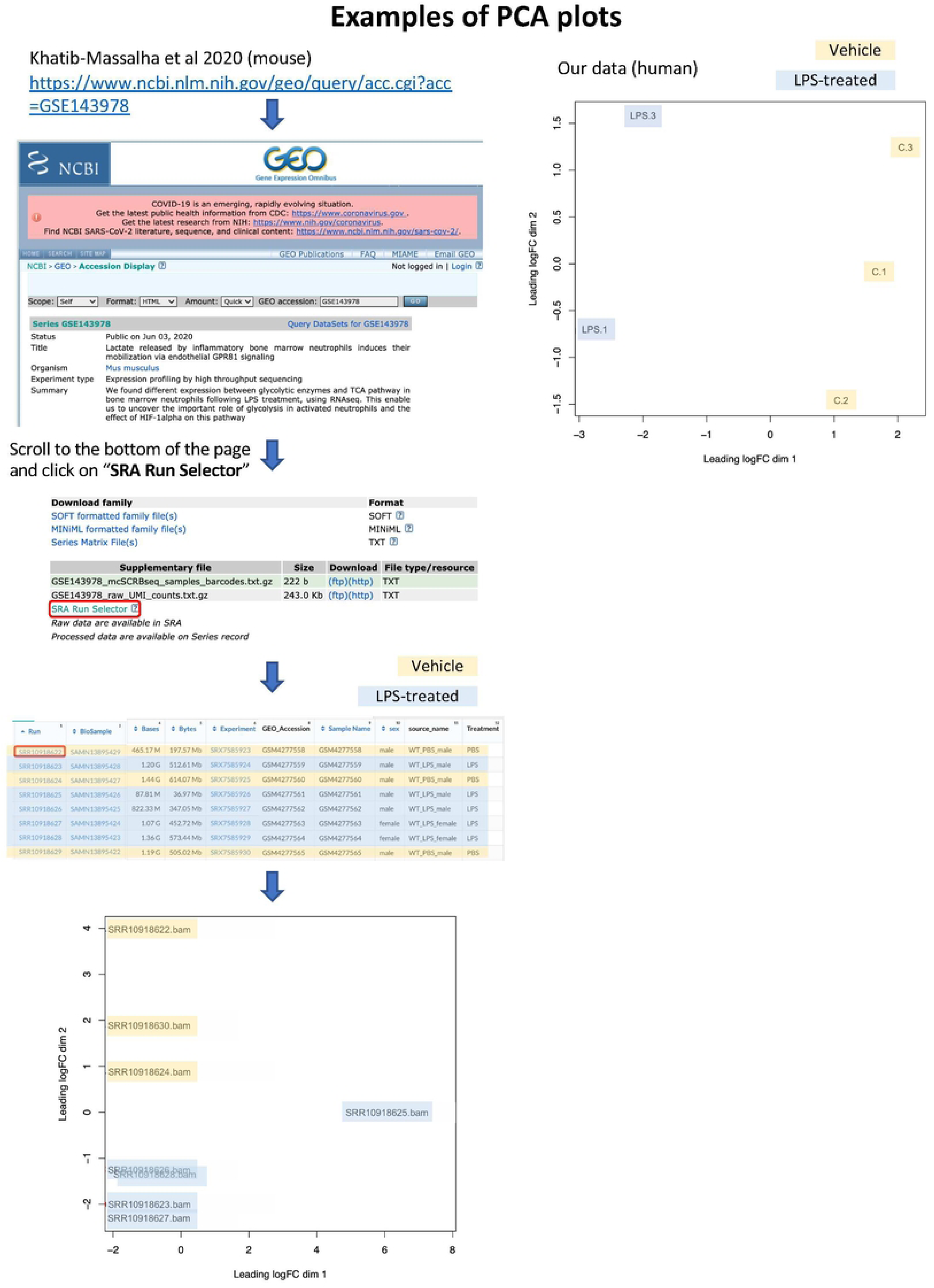
Example of PCA plot in mouse and our human datasets. Similarity between replicates in each group is confirmed by PCA analysis.

**S4 Fig.**
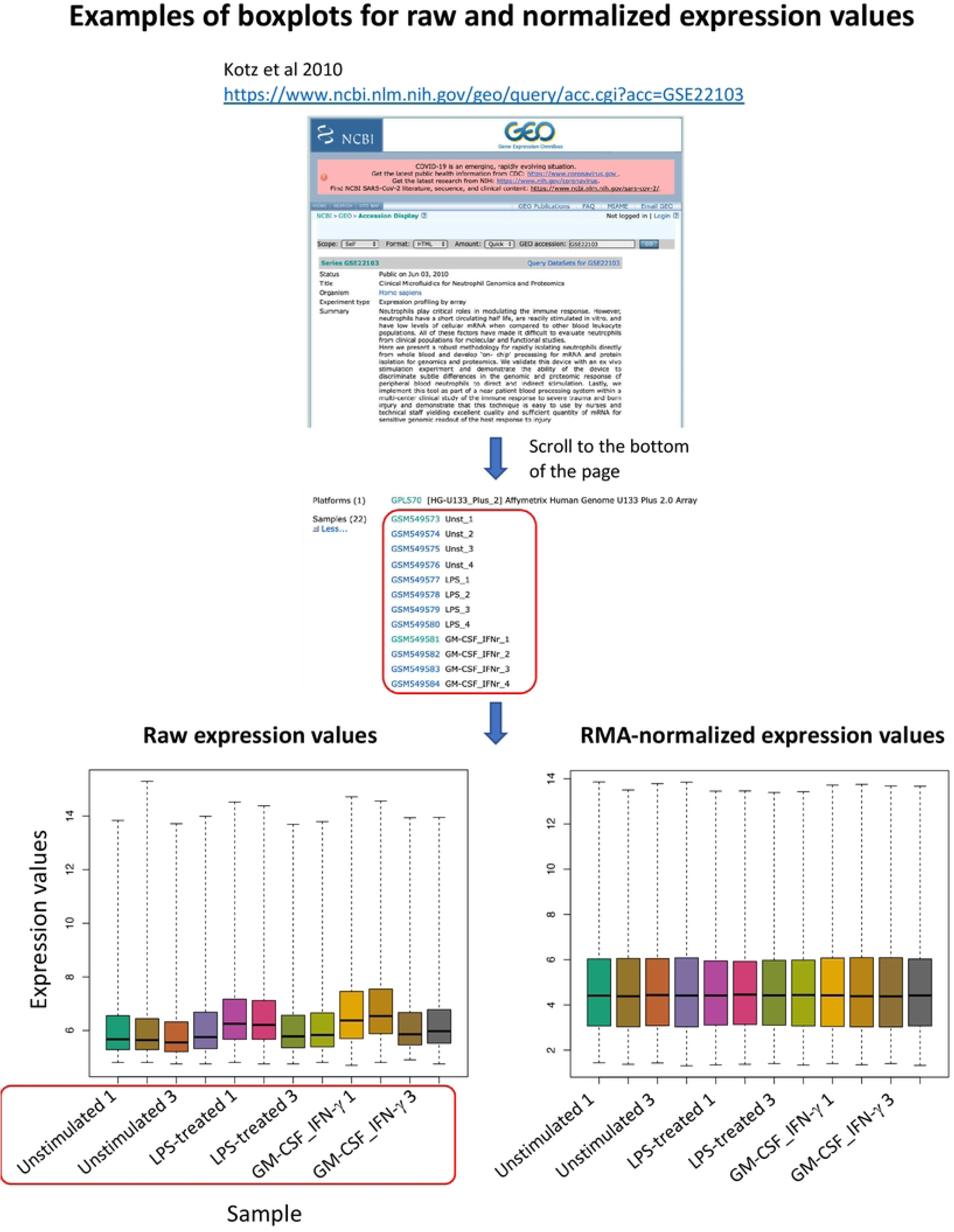
Examples of boxplots for raw and normalized expression values in human microarray datasets. To identify potential outliers, signal distribution of the raw and normalized data was viewed using boxplots. Samples were comparable following background correction and RMA normalization.

**S5 Fig.**
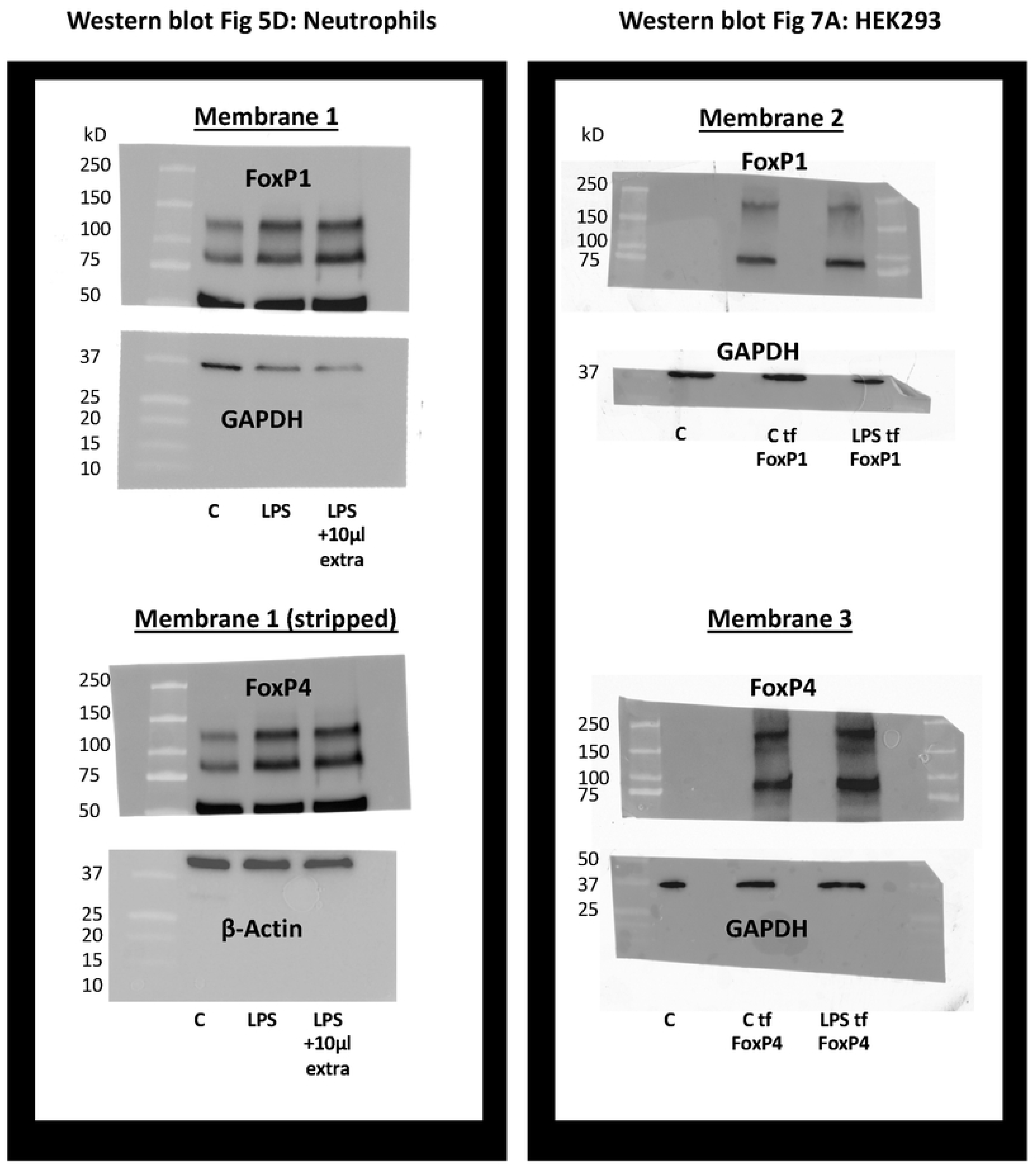
Full western blots of FoxP1 and FoxP4 probed neutrophils and HEK293 cells. The middle band (∼75kda) corresponds to the full-length FoxP1 and FoxP4 protein as described in (22, 63). Tf = transfected HEK293 cells.

**S6 Fig.**
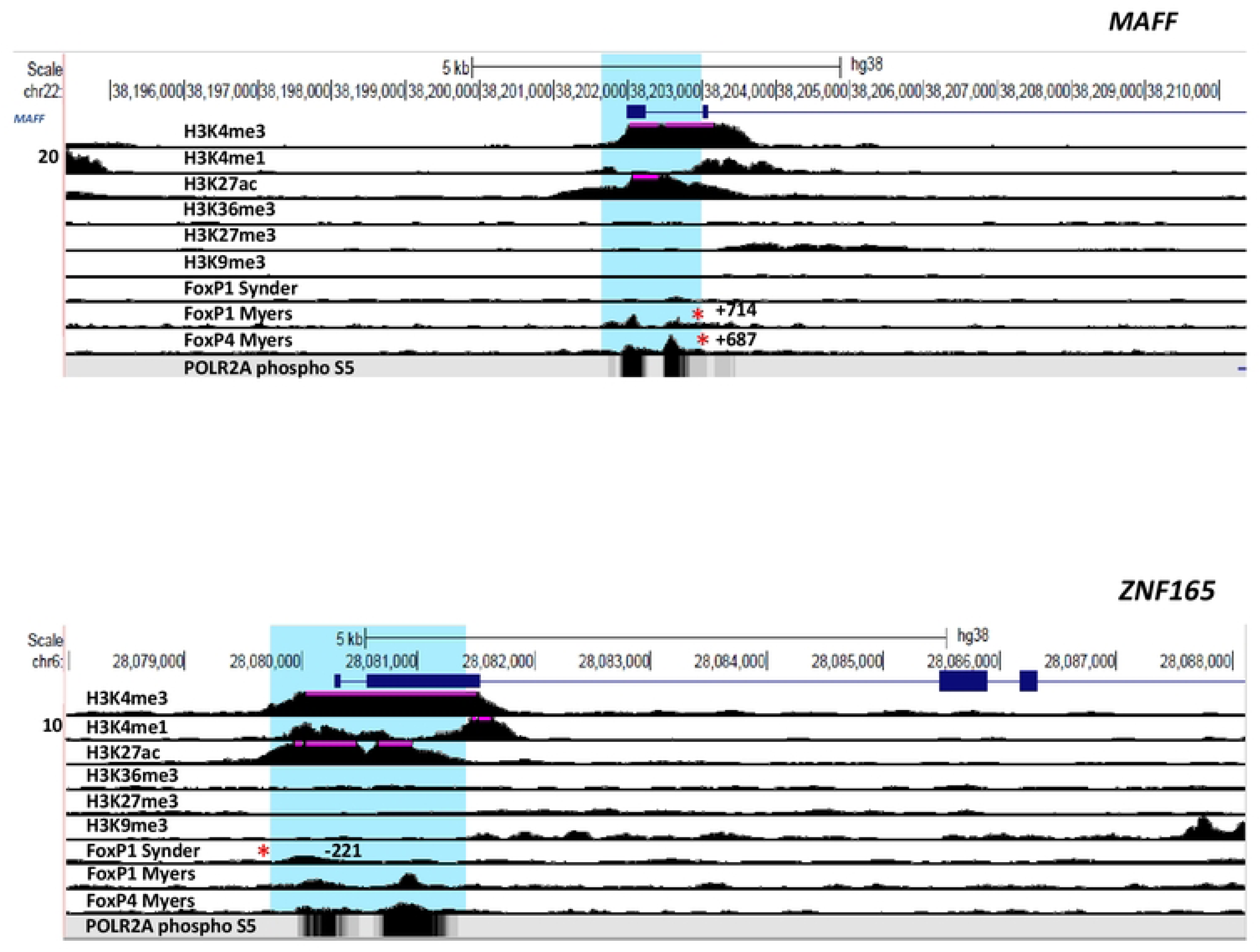
UCSC browser images of FOXP1 and FOXP4 ChIPseq tracks for the *MAFF* and *ZNF165* loci. The areas surrounding the FOXP1 and/or FOXP4 binding sites of genes are shown in blue. Red asterisks represent ChIPseq peaks that contain a consensus motif. Binding sites correspond to regions of enhancer function in neutrophils.

**S7 Fig.**
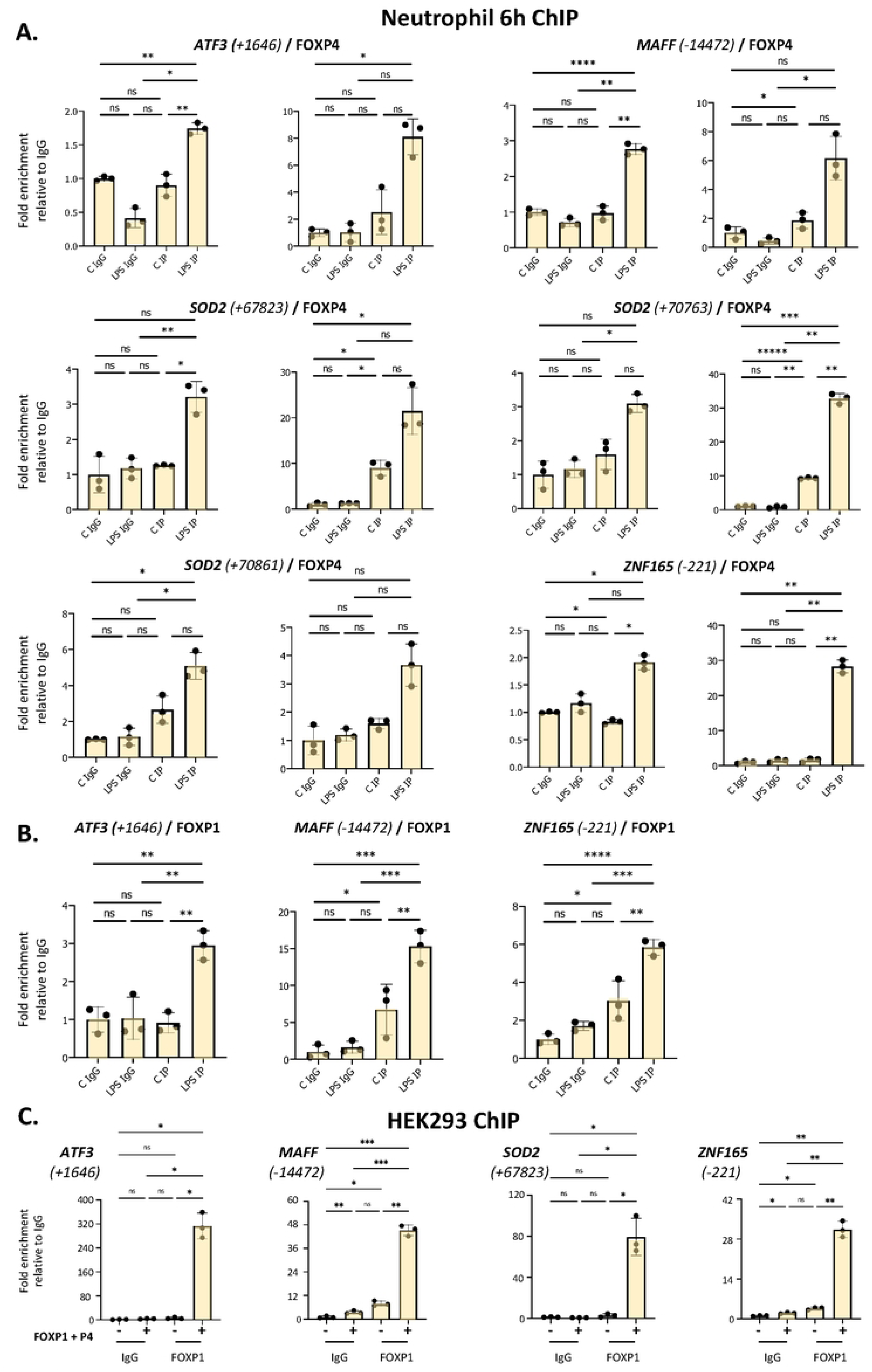
Enhanced binding of FOXP4 and FOXP1 to network gene motifs in individual isolates of LPS-challenged neutrophils and HEK293 cells. **A.** Analysis of the association of FOXP4 with up- and downstream regulatory regions of ATF3, SOD2, MAFF and ZNF165 by ChIP assay in individual isolates of neutrophils treated with or without LPS for 6h (not including the representative data). **B.** Analysis of the association of FOXP1 with up- and downstream regulatory regions of ATF3, SOD2, MAFF and ZNF165 by ChIP assay in one isolate of neutrophils treated with or without LPS for 6h. **C.** Analysis of the association of FOXP1 with up- and downstream regulatory regions of *ATF3*, *MAFF*, *SOD2* and *ZNF165* by ChIP assay in HEK293 cells transfected with and without FOXP1 and FOXP4 expression vectors. Data representative of 2 or 3 biological replicates. All graphics are mean ± SD from 3 technical replicates from a representative sample.*P ≤ 0.05, **P ≤ 0.01, ***P ≤ 0.001, ****P ≤ 0.0001 and ns ≥0.05 as assessed by one-way ANOVAs followed by Tukey’s post hoc test for multiple comparisons. ChIP values are normalized to input for each condition and expressed as a fold relative to non-specific IgG control.

**S8 Fig.**
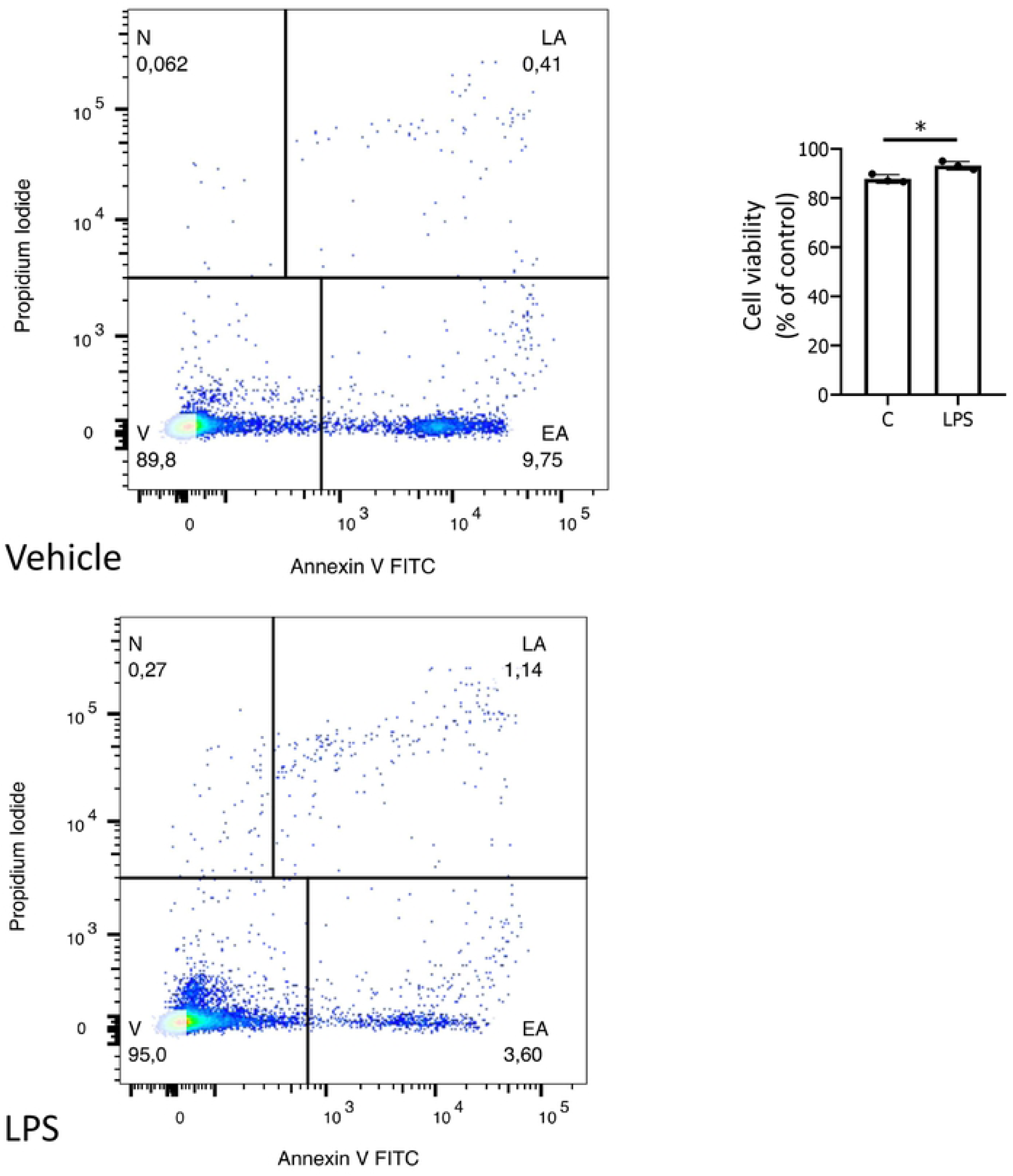
Viability of neutrophils treated for 6h with vehicle and LPS. Cells were gated for singlets, and debris as well as erythrocytes were excluded using the forward and side scatter plot. Viable neutrophils are identified as being negative for both annexin V and propidium iodide. N = non-viable, LA = late-apoptotic, EA = early-apoptotic, V = viable. The data shown is representative of 2 independent experiments. *P ≤ 0.05 by Student’s t-test.

**S1 Table.**
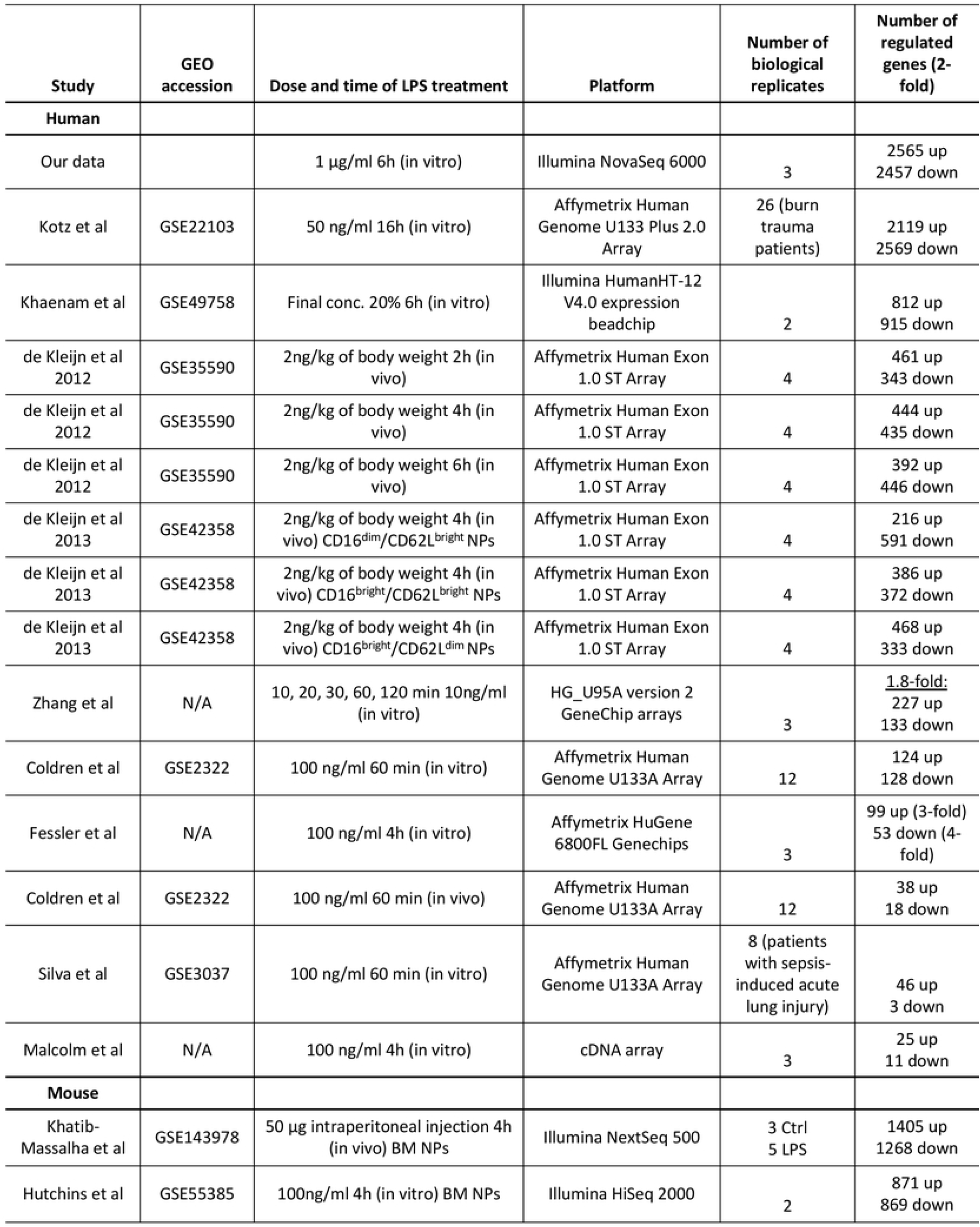
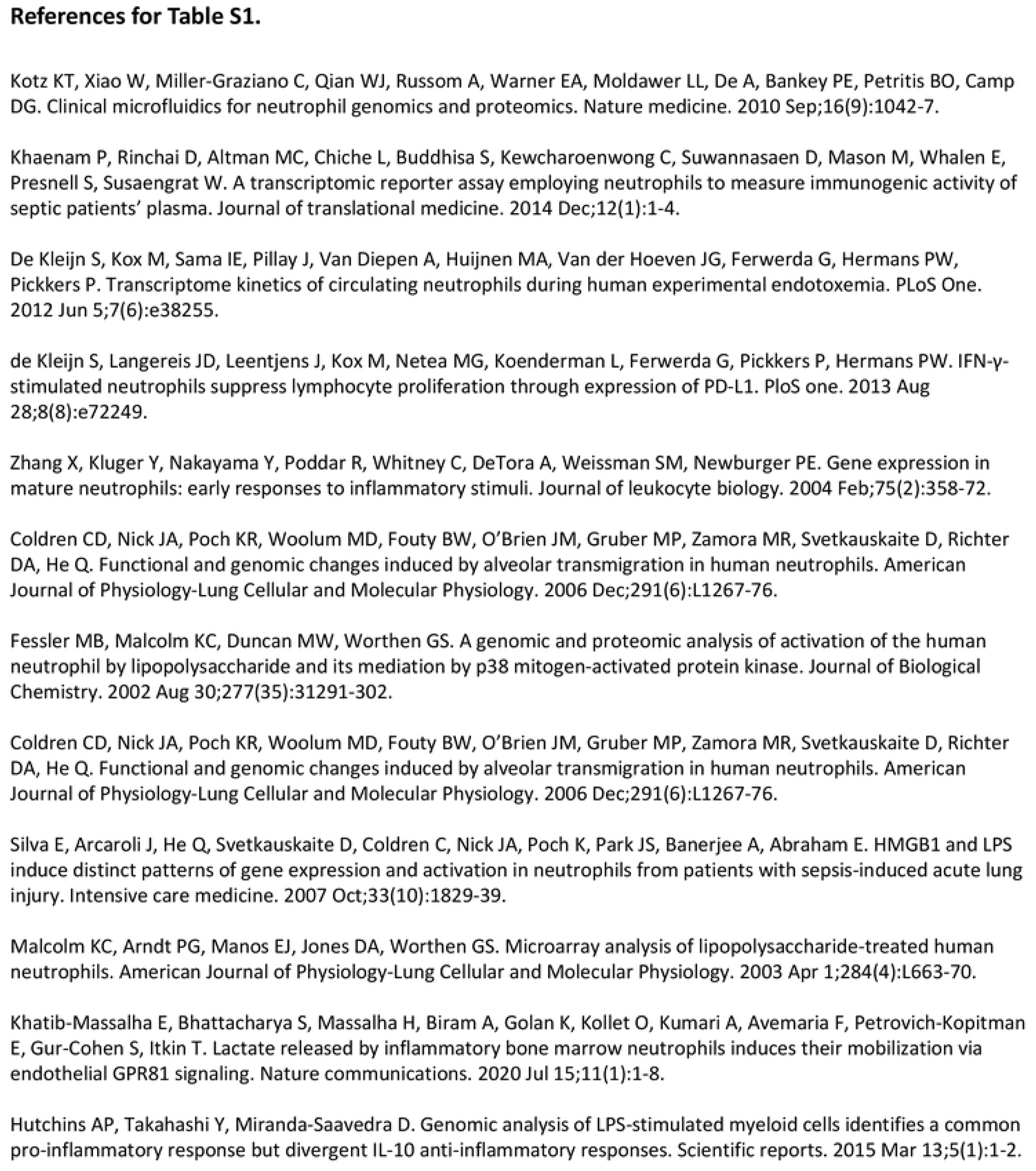
Datasets included in the meta-analysis for LPS-stimulated neutrophils. All datasets included in the analysis with associated publication reference, GEO accession number, dose and time of LPS treatment, platform, and number of biological replicates and 2-fold regulated genes. References are listed on the next page. BM NPs = bone marrow neutrophils.

**S2 Table.** ChIPseq datasets included in this study. All datasets included in the study with associated ENCODE experiment ID (if applicable), target, platform, assembly, lab or publication reference and cell type.

| ENCODE<br>ChIPseq<br>experiment | Target | Platform | Assembly | Lab / Reference | Cell Type |
| --- | --- | --- | --- | --- | --- |
| N/A | FOXP1 | SOLiD/AB<br>sequencer | hg19 | van Boxtel et al 2013 | Colon cancer cell line<br>(DLD1) |
| N/A | FOXP1 | Illumina<br>Genome<br>Analyzer II | hg19 | Gabut et al 2011 | Embryonic stem cells |
| ENCSR029LBT | FOXP1 | Illumina<br>HiSeq 2000 | hg38, hg19 | Michael Snyder Lab | HepG2 |
| ENCSR369YUK | FOXP1 | Illumina<br>NextSeq 500 | hg38, hg19 | Richard Myers HAIB | HepG2 genetically<br>modified with CRISPR |
| ENCSR232LLP | FOXP4 | Illumina<br>NextSeq 500 | hg38 | Richard Myers HAIB | HepG2 genetically<br>modified with CRISPR |
| ENCSR393SYU -<br>ENCFF628GHA | H3K4me3 | Illumina<br>HiSeq 2500 | hg38, hg19 | Bradley Bernstein,<br>Broad | Homo sapiens neutrophil |
| ENCSR586POT -<br>ENCFF897XDW | H3K4me1 | Illumina<br>Genome<br>Analyzer IIx | hg38, hg19 | Bradley Bernstein,<br>Broad | Homo sapiens neutrophil<br>male |
| ENCSR267YXV -<br>ENCFF909JRL | H3K27ac | Illumina<br>HiSeq 2500 | hg38, hg19 | Bradley Bernstein,<br>Broad | Homo sapiens neutrophil |
| ENCSR373WCB -<br>ENCFF356ZPT | H3K36me3 | Illumina<br>HiSeq 2500 | hg38, hg19 | Bradley Bernstein,<br>Broad | Homo sapiens neutrophil |
| ENCSR058FCG -<br>ENCFF981CZJ | H3K27me3 | Illumina<br>HiSeq 2500 | hg38, hg19 | Bradley Bernstein,<br>Broad | Homo sapiens neutrophil |
| ENCSR437MHW<br>- ENCFF777GYB | H3K9me3 | Illumina<br>Genome<br>Analyzer IIx | hg38, hg19 | Bradley Bernstein,<br>Broad | Homo sapiens neutrophil<br>male |
| ENCSR960ALP -<br>ENCFF852BHO | POLR2A<br>phosphoS5 | Not<br>specified | hg38 | Bill Noble, UW | Homo sapiens neutrophil<br>male |
| ENCSR208XPO -<br>ENCFF514ZFZ | POLR2A<br>phosphoS5 | Not<br>specified | hg38 | Bill Noble, UW | Homo sapiens neutrophil |

**S3 Table.** Datasets included in the meta-analysis for neutrophils challenged with various inflammatory signals. All datasets included in the analysis with associated publication reference, GEO accession number, dose and time of LPS treatment, platform, and number of biological replicates and 2-fold regulated genes.

| Study | GEO accession | Dose and time of treatment | Platform | Number of biological replicates | Number of regulated genes (2-fold) |
| --- | --- | --- | --- | --- | --- |
| Human |  |  |  |  |  |
| Our data |  | 1 µg/ml 6h (in vitro) LPS | Illumina NovaSeq 6000 | 3 | 2565 up<br>2457 down |
| Prince et al 2017 | GSE94923 | 1 mM 4h (in vitro) N6/8-AHA (PKA agonist) | Affymetrix Human Genome U133 Plus 2.0 Array | 3 | 1241 up<br>1879 down |
| Kotz et al 2010 | GSE22103 | 20 ng/mL GM-CSF and 100 IU/ml IFN-γ 16h | Affymetrix Human Genome U133 Plus 2.0 Array | 4 | 2799 up<br>2271 down |
| Greenberg et al 2015 | GSE55849 | *Granulibacter bethesdensis* 24h | Affymetrix Human Genome U133 Plus 2.0 Array | 4 | 1791 up<br>1902 down |
| Wright et al 2010 | GSE20151 | *Fusobacterium nucleatum* MOI 1:300 3h | Affymetrix Human Genome U133A Array | 2 | 249 up<br>41 down |

References for Table S3.
Prince LR, Prosseda SD, Higgins K, Carling J, Prestwich EC, Ogryzko NV, Rahman A, Basran A, Falciani F, Taylor P, Renshaw SA. NR4A orphan nuclear receptor family members, NR4A2 and NR4A3, regulate neutrophil number and survival. *Blood*, The Journal of the American Society of Hematology. 2017 Aug 24;130(8):1014-25.
Kotz KT, Xiao W, Miller-Graziano C, Qian WJ, Russom A, Warner EA, Moldawer LL, De A, Bankey PE, Petritis BO, Camp DG. Clinical microfluidics for neutrophil genomics and proteomics. *Nature medicine*. 2010 Sep;16(9):1042-7.
Greenberg DE, Sturdevant DE, Marshall-Batty KR, Chu J, Pettinato AM, Virtaneva K, Lane J, Geller BL, Porcella SF, Gallin JI, Holland SM. Simultaneous host-pathogen transcriptome analysis during *Granulibacter bethesdensis* infection of neutrophils from healthy subjects and patients with chronic granulomatous disease. *Infection and immunity*. 2015 Nov;83(11):4277-92.
Wright HJ, Chapple IL, Matthews JB, Cooper PR. *Fusobacterium nucleatum* regulation of neutrophil transcription. *Journal of periodontal research*. 2011 Feb;46(1):1-2.

**S4 Table.**
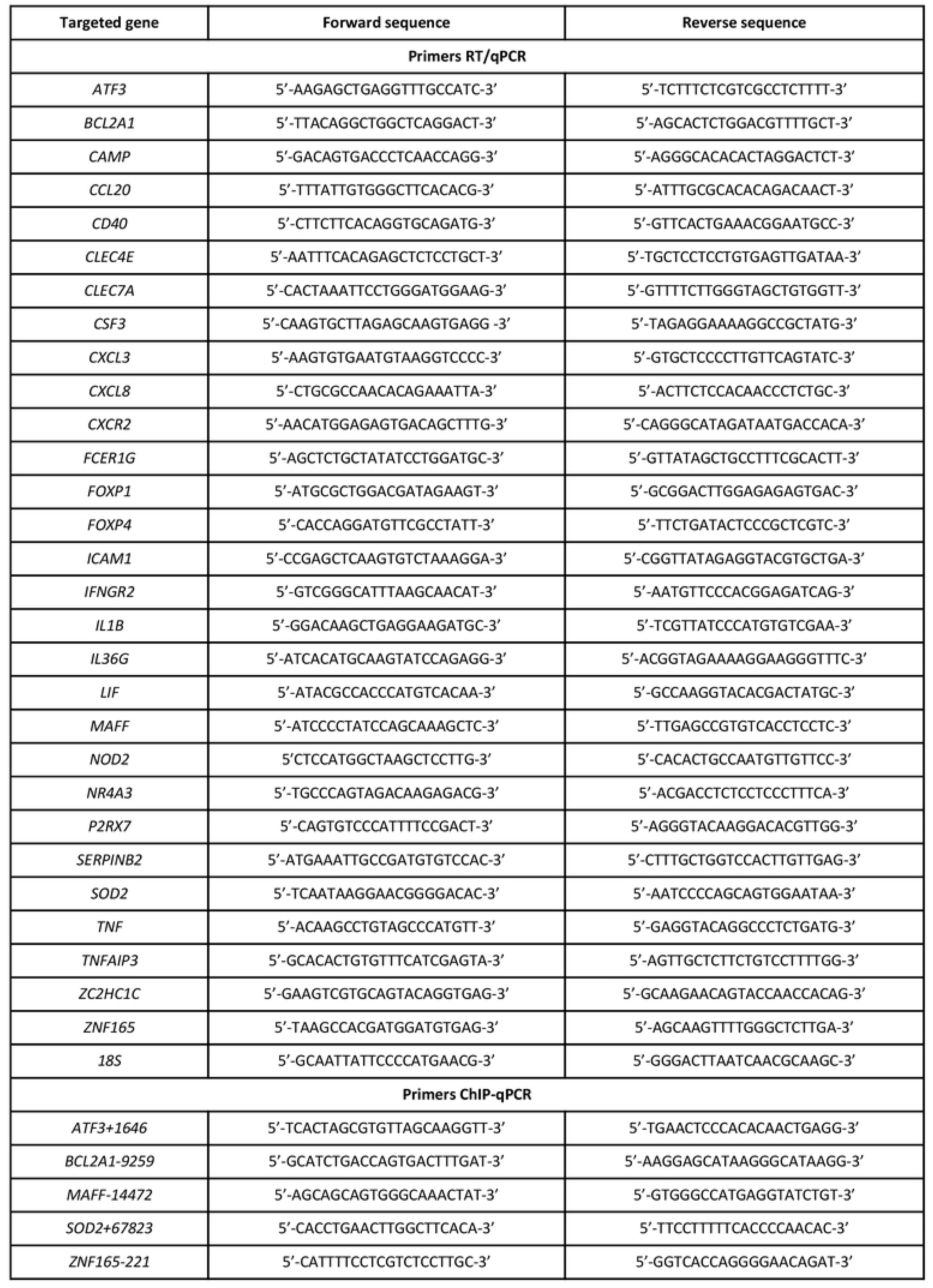
Primer sequences for RT/qPCR and ChIP-qPCR.

**S1 File. RNAseq of primary human neutrophils treated with and without LPS for 6h**. Gene expression changes in LPS and vehicle treated neutrophils identified by RNAseq. Fold change values, along with their corresponding p values, are indicated respectively.

**S2 File. Enriched FOX transcription factor consensus and near-consensus motifs at FOXP1/4 binding sites adjacent to network genes.** Presence and absence of neutrophil histone marks are indicated at each site, represented by + and –, respectively. Bound sites containing a near-consensus sequence (1-2 basepairs mismatch) are highlighted in pink and sites with a consensus motif are highlighted in yellow. Distance to transcription start site (Dist. to TSS) is calculated from the ChIPseq peak center to the TSS of a gene. Each site was statistically significant as assessed by p values less than 0.05 generated from TRAP tools (numbers in light blue; p values combined using Fisher’s method and corrected for multiple testing).

